# ORC1 binds to *cis*-transcribed RNAs for efficient activation of replication origins

**DOI:** 10.1101/2023.03.07.531515

**Authors:** Aina Maria Mas, Enrique Goñi, Igor Ruiz de los Mozos, Aida Arcas, Luisa Statello, Jovanna Gonzalez, Lorea Blázquez, Wei Ting Chelsea Lee, Dipika Gupta, Álvaro Sejas, Shoko Hoshina, Alexandros Armaos, Gian Gaetano Tartaglia, Shou Waga, Jernej Ule, Eli Rothenberg, María Gómez, Maite Huarte

**Author notes:** These authors contributed equally to this study (Enrique Goñi, Igor Ruiz de los Mozos, Aida Arcas).

## Abstract

Cells must coordinate the activation of thousands of replication origins dispersed throughout their genome. Active transcription is known to favor the formation of mammalian origins, although the role that RNA plays in this process remains unclear. We show that the ORC1 subunit of the human Origin Recognition Complex interacts with RNAs transcribed from genes with origins in their transcription start sites (TSSs), displaying a positive correlation between RNA binding and origin activity. RNA depletion, or the use of ORC1 RNA-binding mutant, result in inefficient activation of proximal origins, linked to impaired ORC1 chromatin release. ORC1 RNA binding activity resides in its intrinsically disordered region, involved in intra- and inter-molecular interactions, regulation by phosphorylation, and phase-separation. We show that RNA binding favors ORC1 chromatin release, by regulating its phosphorylation and subsequent degradation. We propose that fluctuating concentrations of RNA during the cell cycle may play a sequential role in controlling origins through interaction with this flexible region of ORC1. Our results unveil a novel non-coding function of RNA as a dynamic component of the chromatin, orchestrating the activation of replication origins.

**One sentence summary:** The human origin recognition complex subunit 1 ORC1, binds to RNAs transcribed from genes with origins of replication at the TSS, which is required for optimal origin activation.

---

The initiation of DNA replication involves a sequential assembly and disassembly of protein complexes on genomic DNA, which is tightly controlled along the cell cycle. Initiation occurs at specific sites throughout the genome where the Origin Recognition Complex (ORC) associates in M/G1. ORC is composed of six subunits (ORC1-6), of which ORC1 is the pioneering subunit in the binding to the chromatin. This binding recruits other members of the complex, followed by additional initiation factors (CDC6, CDT1, MCM helicases, CDC45 and CDC7) in a sequential manner, leading to, (i) licensing and (ii) firing of replication origins (Fragkos et al., 2015). Linked to the firing in S phase, some components of the initiation complex are disassembled and targeted for degradation, which is followed by the activation of DNA helicases and loading of replication factors, avoiding DNA re-duplication events in the same cycle (Kara et al., 2015; Méndez et al., 2002). ORC1 stands out among ORC components in its distinct regulation throughout the cell cycle, which is consistent with its crucial function in the initiation of replication and the maintenance of undamaged cell propagation (Hossain et al., 2021; Méndez et al., 2002).

In *S. cerevisiae*, the position of replication origins is defined by ORC recognition of DNA sequence-dependent elements (Marahrens and Stillman, 1992). In contrast, how origins are positioned in mammalian genomes is still an outstanding question, since ORC does not recognize a known consensus DNA sequence (Vashee et al., 2003). Furthermore, only ∼20% of licensed origins in a given cell fire in S phase (Lebofsky et al., 2006), but how this activation is regulated is not fully understood. Multiple chromatin features are known to influence the flexible selection and activation of origins, which, in combination, dictate the probability of stochastic origin activation. These include chromatin accessibility, specific histone marks such as H4K20me2 (Kuo et al., 2012; Tardat et al., 2010) and H2AZ (Long et al., 2020), and the presence of DNA sequences prone to form G-quadruplexes (G4) (Akerman et al., 2020; Besnard et al., 2012; Valton et al., 2014). While the simultaneous replication and transcription of a precise DNA position are strictly incompatible, the most active origins are localized at transcription start sites (TSSs), with origin activity correlating with the level of gene expression (Dellino et al., 2013; Karnani et al., 2010; Langley et al., 2016; Mesner et al., 2011; Sequeira-Mendes et al., 2009; Sugimoto et al., 2018; Valenzuela et al., 2011). Thus, in the entry of S phase, when early replication origins are fired, RNA is produced in close proximity. This raises the possibility that RNAs could influence origin selection or activation. While previous reports have pointed to specific roles for RNA in replication initiation in *X. laevis* shortly after fertilization (Aze et al., 2017) and at Epstein Barr virus OriP (Norseen et al., 2008), whether transcribed RNA plays a general role in the activation of mammalian origins yet remains unknown.

Here, we show that ORC1 interacts with RNAs transcribed at active origins, which is linked to ORC1 dynamic association to the chromatin and optimal origin firing in S phase.

## ORC1 interacts with RNA in vivo

We hypothesized that ORC1, the first subunit of the initiation complex associating to the chromatin, binds to RNA in cells. *In vitro* RNA binding had been described for human ORC1, mapping to a region between amino acids 413 and 511 (Deng et al., 2009; Hoshina et al., 2013), part of an Intrinsically Disordered Region (IDR) and separated from other domains mediating, among other functions, its binding to nucleosomes (Duncker et al., 2009; Hossain et al., 2021) (**Figures 1A** **and S1A**). In line with this hypothesis, stochastic optical reconstruction microscopy (STORM) on the chromatin fraction of G1-synchronized cells detected a significant cross-correlation between ORC1 and EU-labeled RNA, when compared to randomized images, and not detected in the experimental negative control (**Figures 1B** **and S1B**). These results indicate that ORC1 is in very close proximity to RNA *in vivo*.

**Figure 1.**
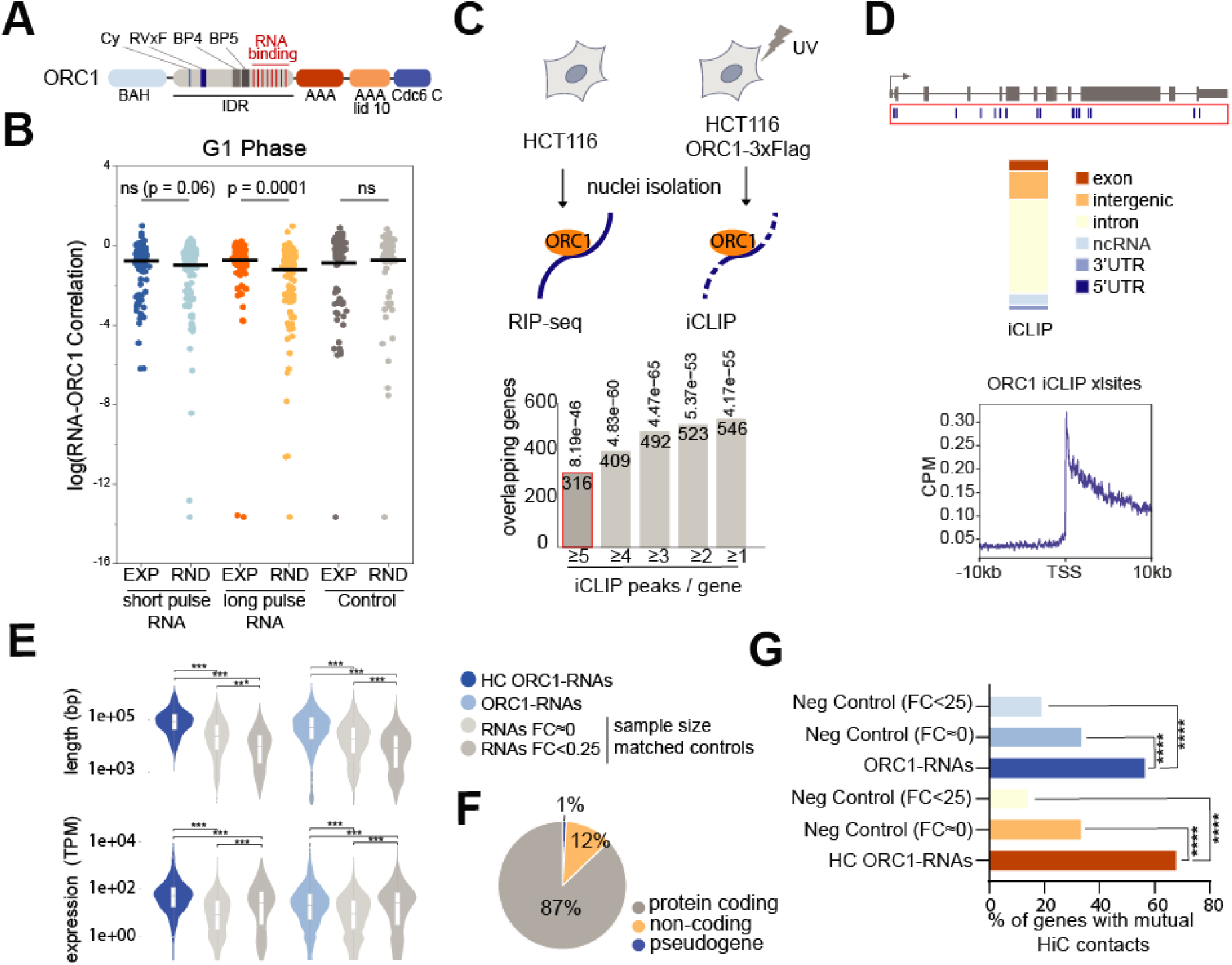
ORC1 binds *in vivo* to RNA. **(A)** Schematic representation of human ORC1 protein domains (El-Gebali et al., 2019; Hoshina et al., 2013; Hossain et al., 2021). **(B)** Mean distribution of cross-correlation between endogenous ORC1, and unlabeled (control) or EU-labeled RNA (short or long pulse), comparing STORM experimental (EXP) and randomized (RND) analysis in the chromatin fraction of U2OS cells synchronized in G1. **(C)** Schematic of RIP-seq and iCLIP experimental approaches, where endogenous or Flag-tagged ORC1 is immunoprecipitated from native or UV-crosslinked nuclear extracts, followed by recovery of full length or digested bound RNAs. Below, number of genes identified by both methods, with different iCLIP stringencies, and hypergeometric test *p-values* of the experimental overlap (RIP-iCLIP) on top of the bars; red for selected high confidence (HC) ORC1-RNAs. **(D)** Genomic distributions of ORC1 iCLIP crosslinks, and their density around TSSs (-/+ 10 kb) of ORC1-bound genes. **(E)** Gene length and expression level of high confidence ORC1-RNAs (HC) and ORC1-RNAs, relative to control genes with different fold changes (FC) in ORC1 RIP-seq. **(F)** Gene biotypes of ORC1-RNAs. **(G)** Percentage of ORC1-RNA and high confidence ORC1-RNA (HC) genes with mutual interactions according to Hi-C analysis, compared to controls shown in Figure 1E.

We then set out to identify RNAs bound by ORC1 by applying complementary approaches (**Figure 1C**). First, we performed native ORC1 RNA immunoprecipitation coupled to sequencing (RIP-seq) of total RNA (ribo-depleted polyA+ and polyA-RNA) from nuclei of dividing HCT116 cells (**Figures S2A and S2B**), using an anti-ORC1 antibody. These experiments identified 2203 RNAs enriched in ORC1 RIP relative to the input samples of nuclear RNA (*Log2 Fold Change*>1, *p-value*<0.05) (**Figure 1C**), while the control IgG did not recover detectable amounts of RNA. To confirm the capacity of immunoprecipitated ORC1 to retrieve RNA, we also performed anti-Flag RIP-seq from nuclear extracts of HCT116 cells transiently expressing ORC1-3xFlag. Most of the identified transcripts (68%) overlapped with the ones interacting with the endogenous ORC1 (Hypergeometric test, *p-value* 1.89e-191) (**Data S1**). Next, theoretical binding predictions were calculated to examine the plausibility of a direct interaction between ORC1 and the co-immunoprecipitated RNAs. *cat*RAPID algorithm, which computes an interaction probability based on the biophysical characteristics of proteins and RNAs (Bellucci et al., 2011), showed that enriched RNAs by RIP also present higher predicted binding compared to the depleted ones (*p-value*<2.2 e^-16^). Moreover, it showed a correlation between theoretical binding score and experimental fold changes in RIP-seq (**Figures S2C and S2D**), thus suggesting that direct binding occurs between the experimentally-defined set of RNAs and ORC1.

We therefore set to detect the direct RNA-protein interactions with nucleotide resolution by applying the iCLIP protocol (Huppertz et al., 2014). HCT116 cells expressing ORC1-3xFlag were UV-crosslinked prior to nuclear isolation and anti-Flag immunoprecipitation. Affinity purified complexes were partially RNase digested, and protected RNAs at and above ORC1-expected molecular weight were extracted, sequenced, and compared with the negative control (i.e. anti-Flag IP in cells that do not express ORC1-3xFlag), which only recovered neglectable RNA amounts (**Figures 1C** **and S2E, and Table S1**). Read alignment to the human genome identified the vast majority of ORC1 crosslink RNA-binding sites (>95%) covering genic regions on the same direction of transcription. While ORC1-RNA crosslinks peaked at the 5’ of genes, they mapped to both exonic and intronic regions, indicating binding to nascent transcripts (**Figures 1D** **and S2F**).

Together, these results demonstrate that ORC1 interacts with RNA in the nucleus of living cells, which could shape the protein functionality along the cell cycle.

## ORC1 interacts with GAA-rich, long, and highly transcribed RNAs, in a sequence-independent manner

Once established that ORC1 interacts with RNA in the nucleus of cells, we next explored whether it prefers to associate through specific RNA motifs. To do that, we searched for motifs in 200nt windows centered around the iCLIP peaks defined with iCOUNT peak caller. This analysis identified the snoRNA C box UGAUGA motif (*e-value* 6.0e-246) (**Figure S2G**), in line with the presence of a high number of crosslinks (35%) corresponding to highly expressed snoRNAs, although only mapping to 6% of the genes of ORC1 bound RNAs (**Data S2**). Consistently, when snoRNAs were filtered out, no motif was found enriched, indicating that the binding of ORC1 to the majority of RNAs (94% of genes) is not mediated through a well-defined sequence. Since ORC1 had been reported to preferentially bind to G-quadruplex RNA structures (Hoshina et al., 2013; Norseen et al., 2009), we also performed G4 predictions around iCLIP peaks, which found a mild enrichment of RNA sequences prone to form this type of secondary structures (**Figure S2H)**. These analyses suggest that ORC1 does not bind to RNA through a specific sequence, although it shows some preference towards structured RNA elements.

To understand the nature of ORC1 RNA interactome, we compared the results of ORC1 RIP-seq and iCLIP experiments. Interestingly, RIP-seq and iCLIP showed good agreement, since ORC1 RNA interactors identified by iCLIP also showed higher fold enrichments by RIP (**Figure S2I**), even when RIP interactors are defined relative to their input, and therefore not biased by expression level. Moreover, by astringently selecting RNAs containing 5 or more iCLIP peaks (1887 genes), a significant overlap (Hypergeometric test, *p-value* 8,19e^-46^) was found with the RNAs identified by RIP-seq (**Figure 1C**), representing high confidence RNAs directly interacting with ORC1 (HC ORC1-RNAs) (**Data S2**). When the selection was extended to genes with more than 4, 3, or 2 iCLIP peaks, the overlap was increased and also highly significant (**Figure 1C**). In addition, several RNA interactors detected by RIP-seq and/or iCLIP were validated in independent RIP and CLIP experiments, with no enrichment detected for the unspecific IgG (**Figures S2J and S2K**), supporting the validity of the used complementary analyses (**Figure S2L**).

We then looked at the characteristics of both the overlapping and the broader set of identified RNAs. HC ORC1-RNAs, as well as the larger set of ORC1-RNAs (**Data S2**), are mostly mRNAs (95% and 87%), with higher expression levels and length than negative control genes (**Figures 1E and 1F**). Similar features were found when assessing RIP-RNAs or iCLIP-RNAs separately (**Figures S3A and S3B**), confirming that ORC1 preferentially binds to RNAs of these characteristics in physiological conditions. According to Repli-seq and Hi-C analyses, the majority of genes encoding for HC ORC1-RNAs and ORC1-RNAs are localized within the early replicating regions of the genome (80%, *p-value* 4e^-253^ and 66.47%, *p-value* 2e^-164^), and interact with each other in the 3D nucleus with higher frequency (*p-value* < 2.2e^-16^ for both sets, **Figure 1G**). Interestingly, sequence analyses performed on full length HC ORC1-RNAs showed the enrichment of tandem GAA repeats (*e-value* 7.5e-^11^, **Figure S3C**), known to be linked to nuclear retention of certain mRNAs (Taniguchi et al., 2007). The sequences were also highly enriched in RNAs detected by RIP-seq of endogenous ORC1 and ORC1-3xFlag, with 82% and 94% of the RNAs containing this type of sequence respectively (**Figures S3D and S3E**). However, the position of iCLIP peaks indicates that ORC1 does not bind RNA through it, as confirmed by the similar EMSA behavior of ORC1-RNAs regardless of the presence of this sequence (**Figures S3F to S3H**). We then concluded that the presence of this motif represents a feature common to many of the RNAs bound by ORC1.

In summary, ORC1 binds to long and highly expressed nuclear RNAs. While this binding appears to be sequence-independent, ORC1-RNA interactors show distinctive features, such as the presence of GAA-repeats and their production from genes that replicate in early S phase and are in 3D proximity.

## RNAs interacting with ORC1 are transcribed from active origins

Since efficient origins are frequently localized near transcription start sites of highly transcribed genes (Dellino et al., 2013; Karnani et al., 2010; Langley et al., 2016; Mesner et al., 2011; Sequeira-Mendes et al., 2009; Sugimoto et al., 2018; Valenzuela et al., 2011), we hypothesized that the binding of ORC1 to RNA could take place in the proximity of the loci where the RNAs are transcribed. We investigated the chromatin at the TSSs of the genes encoding for ORC1-RNAs, by grouping them in 6 quantiles according to their level of ORC1 iCLIP signal, that is, the level of direct ORC1-RNA interaction determined experimentally. This analysis showed a positive correlation between ORC1 binding to RNAs, marks of active chromatin (H3K27ac, H3K4me1, H3K4me3 and H3K36me3), and RNA expression levels, while an opposed trend was observed when considering marks of silent chromatin (H3K27me3 and H3K9me3) (**Figures 2A to 2C, and fig. S4A and S4B**). Interestingly, ORC1 iCLIP signal also correlated with H4K20 methylation, recognized by ORC1 (Kuo et al., 2012), as well as with ORC1 chromatin binding determined by ChIP-seq (**Figures 2D** **and S4C**).

**Figure 2.**
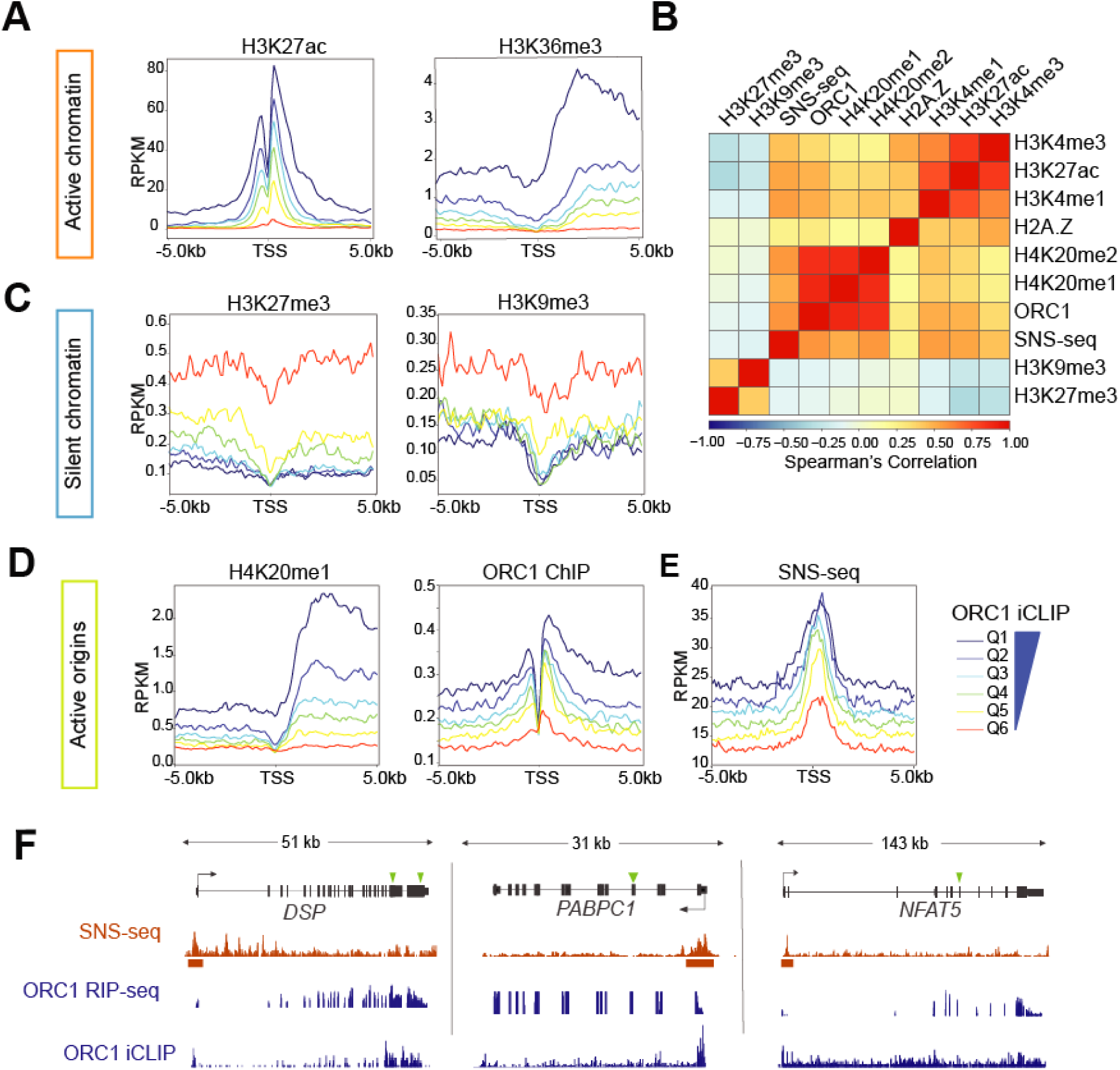
ORC1 binds to RNA produced from active replication origins. **(A)** Density plots of H3K27ac and H3K36me3 ChIP-seq normalized reads across six ORC1 iCLIP-defined quantiles (Q) of ORC1-RNA genes – color legend shown in Figure 2E -, centered around their TSSs (-/+ 5 kb). **(B)** Correlation heatmap between ChIP-seq and SNS-seq data around TSSs of ORC1-RNA genes. **(C, D)** Density plots of **(c)** H3K27me3 and H3K9me3, and **(d)** H4K20me1 and ORC1 ChIP-seq data as shown in Figure 2A (color legend shown in Figure 2E). **(E)** Density plot of SNS-seq normalized reads across six iCLIP-defined quantiles (Q and colored lines) of ORC1-RNA genes, centered around their TSSs (-/+ 5 kb). **(F)** Browser snapshot of representative high confidence ORC1-RNA genes, showing data of ORC1 RNA-binding (ORC1 RIP-seq or iCLIP crosslinks), and replication origins (SNS-seq) at their TSSs, in HCT116 cells. Green arrows indicate positions of GAA repeats.

To further investigate the relationship between ORC1-RNAs and DNA replication, we mapped the active replication origins in HCT116 cells using Short Nascent Strand sequencing (SNS-seq) (**Figure S4D**) (Cayrou et al., 2012). SNS-seq identified 37725 origins that were consistent with previously published origin mapping in other cell types (Akerman et al., 2020), since 63% of the HCT116 origins overlapped with those in quantiles Q1 and Q2 of the 10 defined by the mentioned study, which represent the most robust origins with the highest conservation among cell types (Akerman et al., 2020) (**Figure S4E**). Notably, we found the mapped origins to be highly enriched at the TSS of ORC1-RNA genes (*p-value* 3.82e^-51^, **Figure S4F**). More importantly, the level of ORC1 binding to RNA correlated with the SNS-seq signal, directly proportional to origin activation frequency, as showed by the distribution following the iCLIP gene quantiles (**Figure 2E** **and S4G**).

These results indicate the co-occurrence of ORC1 binding to RNA, and the position and activation of origins in *cis*, that is, at the loci where RNAs are transcribed (**Fig. 2F**).

## RNAs bound by ORC1 are necessary for optimal origin activation

Having determined that ORC1 RNA binding correlates and spatially coincides with origin activation, we addressed whether the RNAs play a role in DNA replication initiation. We first used a global approach by taking advantage of a feature found in many ORC1-RNAs: the presence of several tandem GAA repeats (**Figures S3C to S3E**). We designed antisense oligonucleotides with the sequence TTCTTCTTCTTCTTCTTCTTC (ASO anti-GAA) (**Figure S5A**), which targets RNAs containing tandem GAA repeats for degradation by RNaseH, a strategy previously used to knockdown this type of transcripts (Zheng et al., 2010). RNA-seq showed downregulation of 73% of the RNAs containing a similar GAA motif (Hypergeometric test, *p-value* 1·10^-50^) compared to a scramble control ASO (**Figure S5B**). The knockdown was independently evaluated by RT-qPCR (**Figure S5C**), and RNA-FISH, showing a pattern of foci that were strongly reduced with the transfection of the anti-GAA ASO (**Figure 3A**).

**Figure 3.**
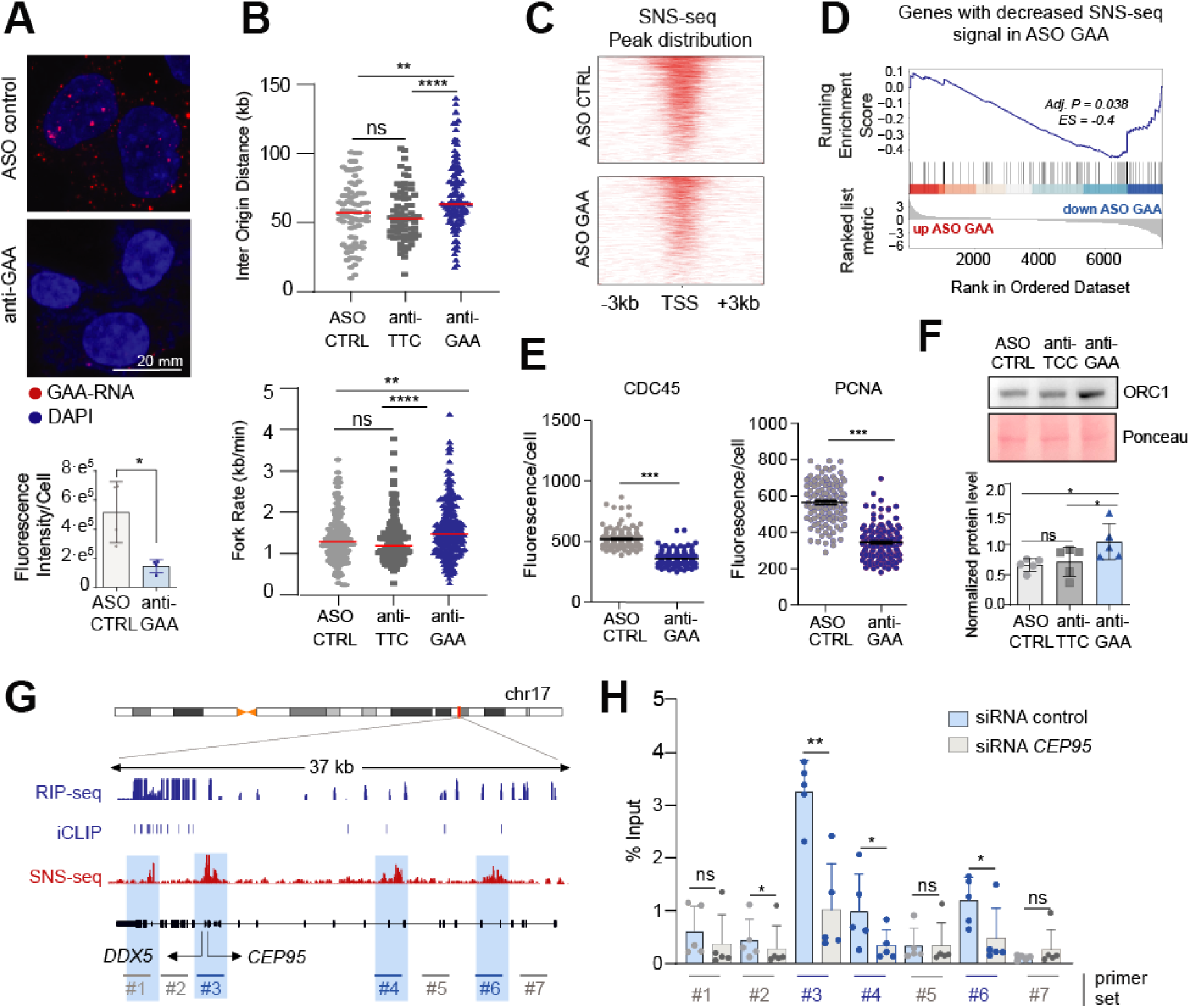
RNAs bound by ORC1 are necessary for optimal origin activation. **(A)** RNA-FISH representative images, and signal quantification (bellow), of RNAs containing GAA repeats, in ASO-transfected HCT116 cells. **(B)** DNA fiber quantification of inter origin distances and fork rates of ASO-transfected HCT116 cells. Red lines indicate median. **(C)** SNS-seq peak count frequency and distribution relative to TSS positions (-/+ 3kb) in ASO control and anti-GAA treated HCT116 cells. **(D)** GSEA showing the reduction of SNS-seq signal (peaks enriched in control condition) in anti-GAA downregulated genes. **(E)** Mean distribution of quantified CDC45 and PCNA chromatin immunofluorescences per cell (HCT116) upon ASO knockdown, after soluble protein washout. **(F)** ORC1 western blot and protein quantification (n>3), in chromatin extracts of ASO-transfected HCT116 cells. **(G)** Browser snapshot at *DDX5*-*CEP95* locus showing ORC1 RIP-seq enrichment, ORC1 iCLIP peaks, and SNS-seq reads in HCT116 cells, with position of qPCR primers (#) indicated, and origins highlighted in blue. **(H)** Enrichment of nascent strands determined by SNS-qPCR at genomic positions indicated in Figure 3G, in siRNA-treated HCT116 cells (n=5).

We then analyzed the effect of GAA-RNAs depletion in DNA replication. To control for effects due to binding of the ASOs to DNA, we also included an ASO targeting the antisense sequence (anti-TTC) (**Figure S5A**). Interestingly, DNA fiber assays showed that the knockdown of GAA-RNAs caused a defect in the activation of replication origins, since increased inter-origin distances and higher fork velocities were detected compared to controls (**Figure 3B**). Indeed, while fewer active origins result in increased distance between origins, the increased fork rate has been described as a mechanism to compensate for origin firing defects (Conti et al., 2007). Consistently, sequencing of nascent strands (SNS-seq) showed a decrease of origin activation at TSSs upon knockdown of GAA-RNAs (**Figure 3C**), suggesting that the RNA may be important for the normal firing of origins at these genomic sites. Of note, the decreased origin activity preferentially affected genes with downregulated RNAs (GSEA *adj. p-value =* 0.038 and *Enrichment Score =* -0.4, **Figure 3D**), pointing to a *cis*-regulatory mechanism of RNA. The detrimental effect on origin activation was also evident when assessing the association to chromatin of the firing factors CDC45 and PCNA, recruiting DNA polymerase to activated origins (Simon et al., 2016), which were significantly reduced on chromatin (**Figures 3E** **and S5D**). In contrast, the levels of ORC1 protein on chromatin were increased (**Figures 3F** **and S5E**). These data support that the general inhibition of RNAs bound by ORC1 has a negative effect on origin activation, linked to an increased association of ORC1 to the chromatin. Nevertheless, we cannot formally exclude that this could be partially attributed to proteins encoded by some of the downregulated mRNAs.

To assess the role of RNA in DNA replication initiation with higher specificity and resolution, we analyzed individual gene loci that produce ORC1-RNAs and contain origins of replication. We selected the *CEP95* locus, transcribing *CEP95* mRNA, one of the HC-ORC1 RNAs. *CEP95* locus contains an origin at the TSS and two downstream secondary origins, according to SNS-seq signal in HCT116 cells (**Figure 3G**). To specifically address the role of *CEP95* mRNA in origin activation, we depleted it using siRNA, since RNAi can downregulate nuclear RNA without interfering with transcription or chromatin structure (Filipowicz et al., 2005) (**Figures S6A to S6C**). Then, we quantified origin activity by SNS-qPCR. Interestingly, the depletion of *CEP95* mRNA caused a specific decrease in the activity of origins on and downstream of *CEP95* TSS, not affecting the origin on the upstream neighbor gene *DDX5* (**Figure 3H**). The deleterious effect of *CEP95* knockdown on local origin activation was also observed by ChIP of CDC45 and PCNA, showing decreased chromatin enrichments at local origins while distant origins were unaffected (**Figures S6D to S6G**). In contrast, *CEP95* overexpression from a plasmid (i.e. *in trans*) did not cause changes in origin activity (**Figures S6D to S6G**), in agreement with a model where transcribing RNAs would be regulating the activity of replication origins found at their site of production. Analogous initiation defects *in cis* were observed when the HC-ORC1 RNA *HSP90AA1* was individually depleted (**Figures S6A to S6C**). The decrease of local origin activity, visualized in the enrichment of nascent strands, was evident, while no consistent effect was appreciated on distant origins at non-related loci (**Figures S6H to S6J**).

## ORC1 RNA binding mutant is impaired in origin activation

To determine how the capacity to bind to RNA is relevant to ORC1 function, we next studied an RNA binding mutant (**Figure 4A**). Based on a previous work (Hoshina et al., 2013), and as evidenced by RNA electrophoretic mobility shift assays (**Figures 4B** **and fig. S7A and S7B**), the substitution to Alanines of three Arginines inside the ORC1 RNA binding region (R441A, R444A and R465A) (**Figure 4A**) results in the loss of *in vitro* RNA binding. The impaired recognition of ORC1-RNAs in cells by the mutant protein was also predicted by *Clever Suite* and *cat*RAPID omics *v2* (**Figures S7C and S7D**), while it did not predict affection of ORC1 ability to bind to DNA (**Figure S7C**). Importantly, the absence of UV-crosslinked RNA in iCLIP experiments (**Figure S7E**) and the decreased co-localization with RNA by STORM (**Figure 4C**) showed experimentally the lack of RNA binding by the mutant ORC1 in cells.

**Figure 4.**
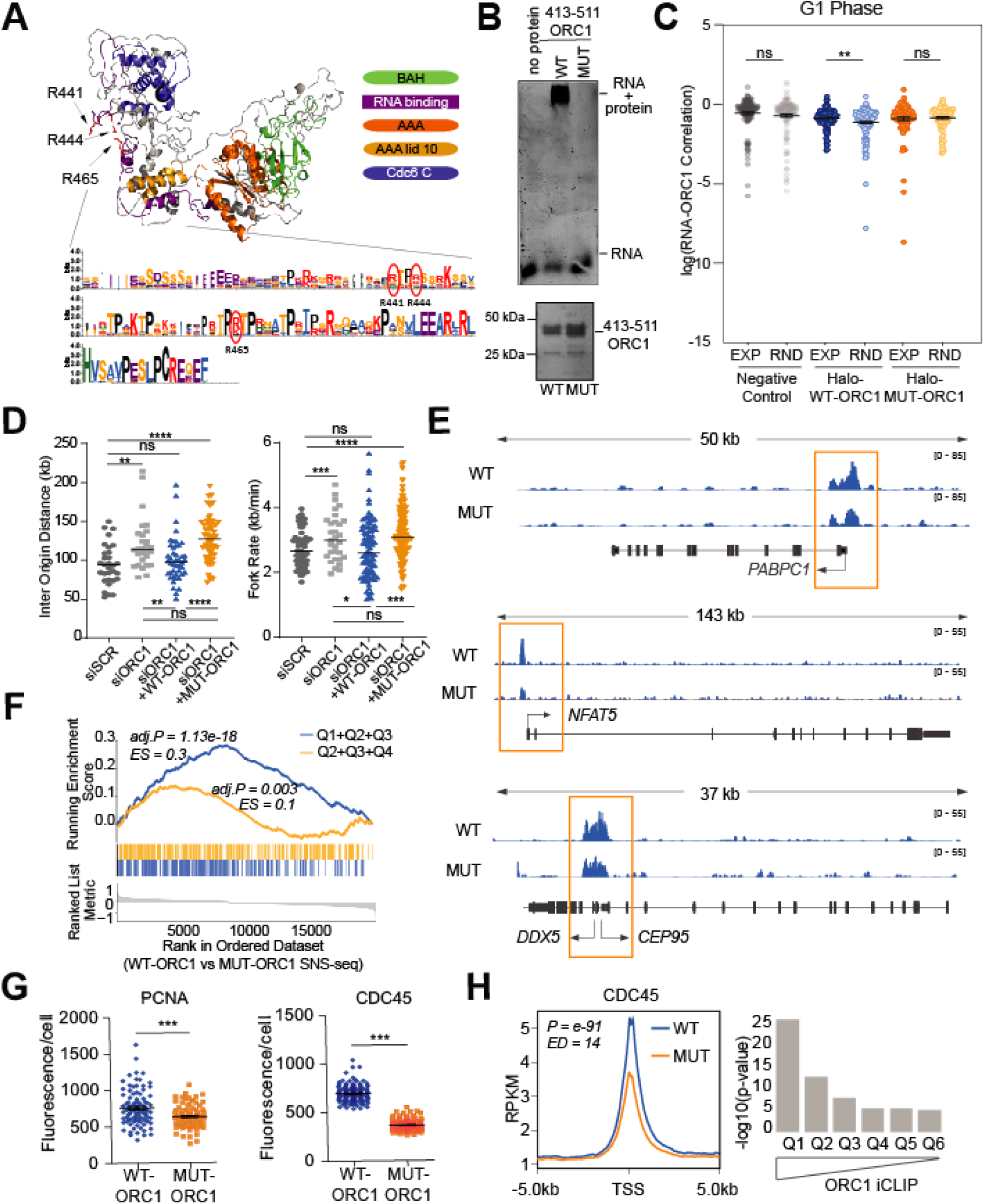
ORC1 RNA-binding mutant is impaired in origin activation. **(A)** 3D model of human ORC1 showing domains in colors, and residues R441, R444 and R465 (involved in RNA-binding) in red. Below, vertebrate consensus of ORC1 RNA-binding region, circles indicating mutated residues in MUT-ORC1. **(B)** RNA staining of EMSA assays, with GST-tagged purified WT and MUT-ORC1 (amino acids 413-511) (2.5 µM) incubated with fragmented cellular RNA (2.5 µM). Below, silver staining of proteins used in the assay. **(C)** Mean distribution of cross-correlation between ORC1 and EU-labeled RNA (long pulse) in G1-synchronized U2OS cells, untransfected or transfected with Halo-tagged WT and MUT-ORC1, comparing STORM experimental (EXP) and randomized (RND) samples. **(D)** DNA fiber quantification of inter origin distances and fork rates in HCT116 cells transfected with the indicated siRNAs, +/- plasmids expressing Flag-tagged WT or MUT-ORC1. Black lines indicate median. **(E)** Browser snapshot at ORC1-RNA *PABPC1*, *NFAT5* and *DDX5-CEP95* loci, showing SNS-seq normalized signal of HCT116 cells stably expressing WT or MUT-ORC1. **(F)** GSEA showing enrichment of ORC1-RNAs in merged iCLIP-defined quantiles (Q), towards ranked genes according to their WT vs MUT (*log2 Fold Change*) SNS-seq coverage at TSSs. **(G)** Mean distribution of quantified CDC45 and PCNA chromatin immunofluorescences per cell, in HCT116 cells stably expressing WT or MUT-ORC1, after soluble protein washout. **(H)** Coverage plot of CDC45 ChIP-seq data at TSSs, in WT or MUT-ORC1 HCT116 stably expressing cells, and *t*-test statistical results between their coverage at TSSs of ORC1 iCLIP-defined gene quantiles (Q).

We next investigated whether the usage of ORC1 RNA binding mutant could affect origin firing. Cells depleted of the endogenous ORC1 and transfected with plasmids expressing WT or MUT-ORC1 were subjected to fiber assays. As expected, depletion of the endogenous ORC1 resulted in decreased number of fired origins, reflected by increased distances between origins (Coulombe et al., 2019) and compensatory increased fork speed (**Figure 4D, and fig. S7F and S7G**). While WT-ORC1 rescued the normal firing and fork rates, MUT-ORC1 did not (**Figure 4D**), suggesting that the RNA-binding activity of ORC1 is needed for optimal DNA replication initiation. Notably, SNS-seq showed decreased origin activity in cells stably expressing MUT-ORC1, and preferentially affecting origins at genes that produce the ORC1-RNAs with stronger level of binding to ORC1 as determined by iCLIP (GSEA *Adj. p-value=*1.13e-18 for genes within Q1-Q3 iCLIP quantiles vs 0.003 for Q4-Q6, and *Enrichment Scores* of 0.3 and 0.1 respectively, **Figures 4E and 4F, and fig. S7H**). These results support the idea that ORC1 binding to RNA has a *cis*-regulatory effect on replication origins, as observed upon global (**Figures 3D, and fig. S5A to S5C**) or individual (**Figure 3H** **and S6**) RNA knockdown. Also in line with fewer origin activation events observed in GAA-RNA-depleted cells (**Figures 3E** **and S5D**), PCNA and CDC45 were significantly reduced at the chromatin in cells expressing ORC1 RNA-binding mutant (**Figures 4G** **and S7I**), also revealed by ChIP-seq signals, primarily decreased at TSSs of transcribing RNAs bound by ORC1 (**Figure 4H**).

We then concluded that the use of ORC1 RNA-binding mutant recapitulates the origin activation defects observed upon knockdown of RNA interactors of ORC1, suggesting that RNA-binding activity of ORC1 is important for normal initiation of DNA replication.

## RNA binding facilitates ORC1 phosphorylation and chromatin release

To better understand how RNA-binding could be important for ORC1 function, we further investigated the behavior of the RNA-binding mutant by analyzing its chromatin association. Notably, MUT-ORC1 was more associated to the chromatin than its wild type counterpart (**Figure S8A**), as was the endogenous ORC1 when ORC1 RNA interactors were depleted (**Figures 3F** **and S5E**). Indeed, MUT-ORC1 presented longer half-life than WT-ORC1, as observed upon translational inhibition with cycloheximide treatment, while both proteins responded to proteasomal inhibition at a similar rate (**Figure 5A**), suggesting that the RNA binding activity of ORC1 decreases its stability. Importantly, while the knockdown of GAA RNAs also led to increased levels of WT-ORC1 on chromatin, it had no effect on the levels of MUT-ORC1 (**Figure S8B**), indicating that this phenotype is RNA-dependent.

**Figure 5.**
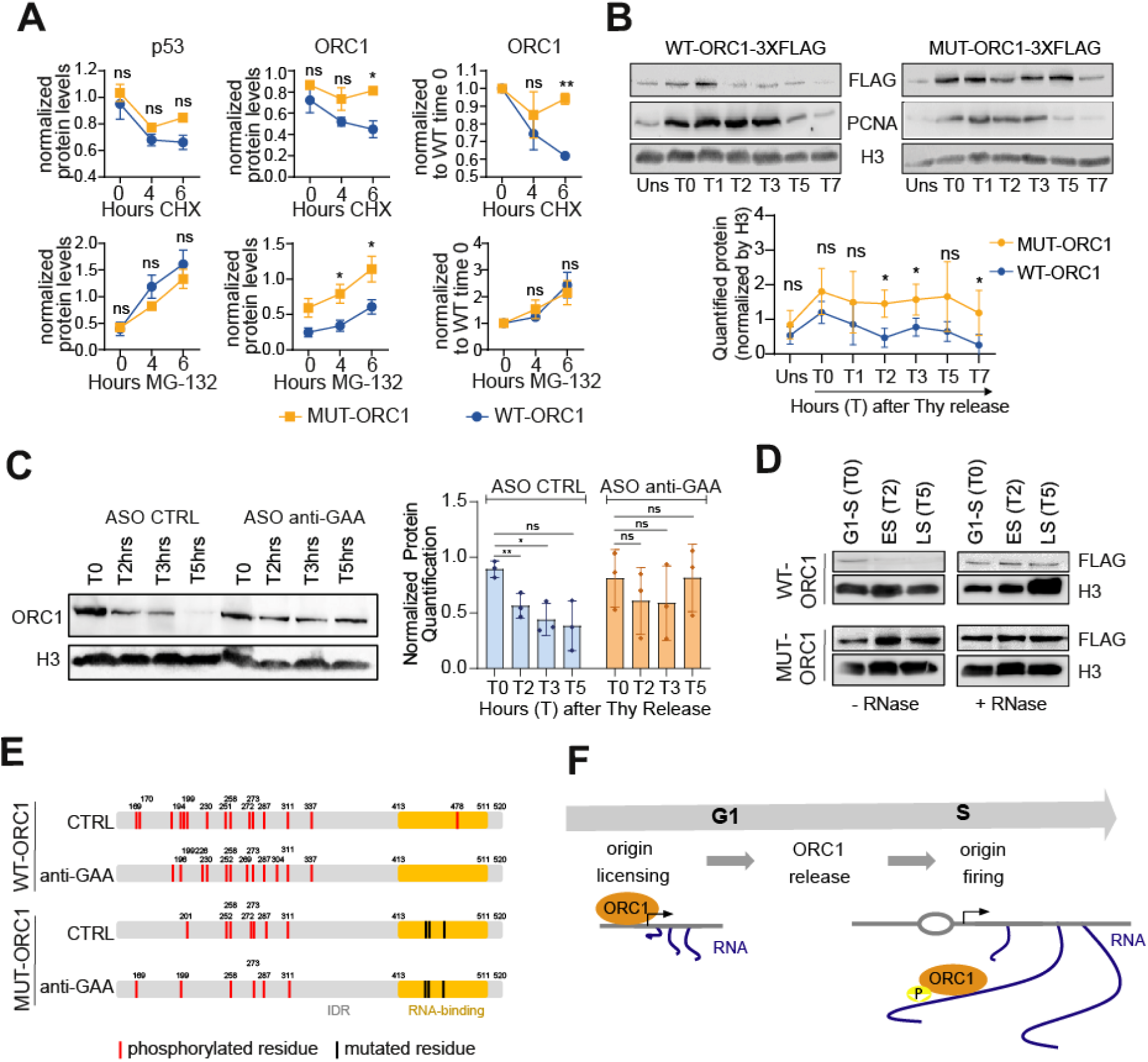
RNA regulates origin activation by controlling ORC1 chromatin release. **(A)** p53 and ORC1-3xFlag protein quantification from western blots with total extracts of HCT116 cells, transfected with WT or MUT-ORC1, and treated with cycloheximide (CHX) or MG-132 (n=3). **(B)** Western blot on chromatin extracts of HCT116 cells, transfected with Flag-tagged WT-ORC1 and MUT-ORC1, unsynchronized (Uns) or synchronized in G1/S and released at different times (T as in *Figure S8C*). Below, normalized protein quantification (n=4). **(C)** Western blot and quantification of endogenous ORC1 on chromatin in different stages of the cell cycle (T as in *Figure S8C*), upon depletion of GAA RNAs (ASO anti-GAA) or control conditions (ASO CTRL). **(D)** Western blot showing the effect of RNase A treatment on WT and MUT-ORC1 chromatin association, along the cell cycle of synchronized cells (T as in *Figure S8C*). **(E)** Representation of the IDR in WT and MUT-ORC1, showing RNA-binding regions (orange), and the discrete positions of RNA-binding mutations (black) and phosphorylated residues (red) detected by mass spectrometry, in control or GAA-knockdown conditions. **(F)** Model for RNA binding to ORC1 in different phases of the cell cycle. After origin licensing, transcribed RNA regulates ORC1 release and protein phosphorylation levels, which is required for proper origin activation.

Since origin licensing and firing is governed by the temporal chromatin association and release of ORC along the cell cycle, we decided to dissect this phenotype in synchronized cells. Even though both WT-and MUT-ORC1-expressing cells progressed through S phase at similar rates (**Figure S8C**), they showed striking differences in ORC1 chromatin association dynamics (**Figure 5B**). WT-ORC1, but not MUT-ORC1, was released from the chromatin at the entry of S phase, suggesting that the observed persistence of MUT-ORC1 on chromatin was due to an inefficient protein release. Importantly, endogenous ORC1 also showed the delayed chromatin release when GAA-RNAs were depleted (**Figure 5C**). Similar effects were observed for WT-ORC1 when synchronized cells were permeabilized and treated with RNase A, resulting in an enhanced chromatin association along S phase that is progressively lost with no RNase A treatment (**Figures 5D** **and S8D**). In contrast, increased MUT-ORC1 chromatin persistence was observed regardless of RNA degradation (**Figures 5D** **and S8D**). Of note, the delay in MUT-ORC1 release was associated with reduced levels of PCNA loaded on chromatin (**Figure 5B**), indicating that the RNA-dependent release of ORC1 is linked to origin activation.

Given that both origin firing and ORC1 stability are governed by phosphorylation (Fragkos et al., 2015), we hypothesized that the phenotypes observed may account for an RNA-dependent modulation of ORC1 phosphorylation. In line with this hypothesis, MUT-ORC1 showed lower mobility in gels, as did the endogenous ORC1 upon the knockdown of GAA-RNA (**Figure S8E**). Phosphoproteomic analysis confirmed the hypo-phosphorylation of MUT-ORC1 (**Data S3**), particularly affecting its IDR, where phosphorylated sites are known to regulate ORC1 chromatin release and proteasomal degradation (Méndez et al., 2002) (**Figure 5E, and fig. S8F and S8G**). Decreased phosphorylation of WT-ORC1 was also observed upon GAA-RNA knockdown, while total phosphorylation levels of ORC1 RNA-binding mutant, as well as the number of phosphorylated residues, were less affected by the transient downregulation of ORC1 RNA interactors (**Figures 5E** **and S8G**), indicating that the interaction with RNA plays a role in regulating ORC1 phosphorylation, and subsequent chromatin release and degradation prior to origin firing. Our results suggest a reciprocal regulation of RNA binding and phosphorylation at the IDR of ORC1, which is linked to its chromatin release and origin activation (**Figure 5F**).

Together, here we demonstrate that RNA binding dynamically modulates ORC1 function by favoring its chromatin release, which is required for efficient origin activation.

## Discussion: RNA as a dynamic regulator of ORC1 function

Here, through orthogonal approaches, we show that RNA, the product of transcription, is needed for optimal origin activation mediated through the intrinsic RNA binding capacity of ORC1. We demonstrate that RNA transcribed in the proximity of the origins influences the efficiency at which they are activated. In line with this model, we find an enrichment of ORC1 iCLIP crosslinks at the 5’ region of the RNAs, although not exclusively restricted to it. This is not unexpected given the described mechanism, and the fact that RNA transcripts remain close to the chromatin while co-transcriptionally processed. Moreover, actively transcribed genes are known to form loops, maintaining 3D proximity between the 5’and 3’ ends (Kuehner et al., 2011). Specifically, we found that the binding of RNA by ORC1 favors efficient release of the protein at the entry of S phase. As suggested in previous studies (Kara et al., 2015), our data indicates that ORC1 release is a pre-requisite for origin firing, although the precise sequence of molecular events at this critical step will require further studies to be fully understood.

Of note, we failed to detect ORC1 RNA binding at a fraction of active origins. While this could be due to limited sensitivity, the RNA-dependent regulation of origins may be restricted to the genes with active transcription, known to replicate in early S phase, where we find the genes producing ORC1-RNAs. Origins found at transcriptional deserts, associated with late replication, might be activated through a mechanism not involving RNA interaction with ORC1. On the other hand, ORC1 might not be the only sensor of RNA at origins from transcriptionally active regions. RNA may interact with additional proteins in the initiation complex. Among them, CDC6 and CDT1 possess positively charged IDRs (Parker et al., 2019) with repetitive sites similar to the observed in ORC1IDR, although their interaction with RNA has not been studied.

Short linear protein motifs (SLiMs) inside ORC1 IDR have been implicated in ORC1 self-interaction as well as interaction with Protein Phosphatase 1α (PP1α) and CDC6 (Hossain et al., 2021), and, regulated by phosphorylation and dephosphorylation, they contribute to the control of ORC1 levels during the cell cycle. Interestingly, the described SLiMs do not overlap with ORC1 RNA binding domain albeit are part of the same highly flexible region (Hossain et al., 2021) (**Figure 1A**). Thus, the interaction of ORC1 with RNA may influence interactions with these factors and/or other properties linked to the IDR. Among those, ORC1 IDR has been implicated in *in vitro* liquid-liquid phase separation enhanced by DNA (Hossain et al., 2021; Parker et al., 2019), being CDK/Cyclin phosphorylation a key inhibitor of liquid phase condensation by replication initiation factors (Parker et al., 2019). While the effect of RNA was not tested in those studies, RNA biophysical properties are consistent with an analogous role. Supporting this notion, we observed that RNA induces the formation of droplets of WT-but not of MUT-ORC1 in a concentration-dependent manner, reaching a threshold where RNA leads to droplet dissolution (**Figures S9A to S9C).**

Based on our data, and by analogy to the RNA feedback mechanism described in transcription (Henninger et al., 2021), where the balance between positive and negative charges determines whether RNA promotes formation or dissolution of IDR-containing protein condensates, RNA could be playing a sequential role in the initiation of replication (**Figure S9D**). We speculate that in late M phase, incipient transcription may favor the formation of coacervates that nucleate ORC1 around TSSs, explaining the presence of ORC1 bound to RNAs transcribed from active origins at early replicating genomic regions, as well as the enrichment of iCLIP binding sites at 5’ ends of genes. Nevertheless, the need of other factors for ORC1 chromatin recruitment should not be excluded. Once preinitiation complexes are assembled on DNA, the property of forming condensates would not be further required. On the contrary, higher local RNA concentration resulting from transcription elongation, particularly at longer genes, would help ORC1 disassembly from chromatin, coupled to an RNA-dependent control of ORC1 phosphorylation balance, and followed by its targeting for degradation and origin firing. This would also explain the enrichment of RNAs containing GAA repeats among ORC1 partners, since these purine-rich sequences are thought to retain RNAs in the nucleus through a saturable nuclear retention factor (Taniguchi et al., 2007).

In addition to the described mechanisms, ORC1 is known to have cellular functions beyond the origins of replication (Popova et al., 2018; Prasanth et al., 2010). It is likely that some of the ORC1-RNA interactions are occurring at those locations. For instance, the interaction between ORC1 and RNA may also be involved in ORC1 transport in and out of the nucleus (Kopytova et al., 2016), which may account for the presence of fully spliced RNAs among ORC1 binders; all questions deserving future investigation.

Overall, our results unveil a novel non-coding function for RNA as a dynamic component of the chromatin, which helps to coordinate transcription and replication in the nucleus.

## Acknowledgements

We thank Dr. Bruce Stillman for the kind gift of anti-ORC1 antibody and ORC1 constructs; Dr. Juan José Lasarte for helping with protein purification; Amaya Abad for NGS library preparation and Dr. Francesco Marchese and Tomás Aragón for critical reading of the manuscript.

This work was supported by grants to M.H.: PID2020-113683GB-100 / funded by MCIN/ AEI /10.13039/501100011033, La Caixa Foundation [LCF/PR/ HR21/00176], European Research Council Consolidator 771425 and Worldwide Cancer Research grants 20-0204; and to G.G.T: European Research Council [RIBOMYLOME_309545 and ASTRA_855923]. A.M.M. was supported by PhD fellowship from Ministerio de Economía y Competitividad REF: BES2015-074569 cofounded by FSE.

## Contributions

A.M.M. designed, performed and analyzed most of the experiments; E.G. performed computational and statistical analyses; I.R.M. performed iCLIP and RIP-seq data analyses and metanalyses; A.A. performed data analyses and protein modeling and predictions; L.S. performed experimental work; J.G. assisted with experimental work.; L.B. performed iCLIP experiments supervised by J.U.; W.T.C.L. and D.G. performed STORM experiments in collaboration with A.M.M. and supervised by E.R.; A.S. assisted with experimental work; S.H. purified recombinant ORC1 protein supervised by S.W.; A. A and G.G.T. performed *catRapid* analyses; M.G. supervised A.M.M. in SNS-seq and DNA fiber assays; M.H. conceived and designed the study, supervised the work, obtained funding and wrote the manuscript with the input from all authors.

## STAR methods

### Cell lines, growth conditions and culture treatments

HCT116 cells were cultured in RPMI-1640 (GIBCO), and U2OS cells in DMEM (GIBCO), all mediums supplemented with 10% fetal bovine serum (GIBCO) and 1x penicillin/streptomycin (Lonza), and maintained at 37°C and 5% CO_2_. To generate HCT116 cells stably expressing WT-ORC1-3xFlag or MUT-ORC1-3xFlag, cells were transfected with pcDNA3.1 vectors containing codon-optimized wild type or mutated cDNA sequences of ORC1 (**Table S2**), and treated with Neomycin-G418 (Sigma) 500 µg/mL for 10 days.

Proteasome inhibition was achieved with short pulses (4 and 6 hours) of MG-132 (MilliporeSigma) 50 µM. Cycloheximide (Sigma) was incubated in culture medium at 100 µg/mL for 4 or 6 hours for translation inhibition.

For RNA *in vivo* labelling, synchronized U2OS cells were incubated with 5-ethynyl uridine (EU-ThermoFisher) at 0.2 mM final concentration in complete culture medium, in short (20 minutes) or long (16 hours and 3 hours release) pulses.

For fiber assays, exponentially growing HCT116 cells were pulsed with 50 µM CldU (Sigma) for 20 minutes, washed, and subjected to a second 20 minutes pulse with 250 µM IdU (Sigma).

In experiments with RNase A treatment, synchronized cells at different points of S phase (see cellular synchronization section) were trypsinized, permeabilized with 0.05% Tween-20 in PBS for 10 min on ice, and mock-treated or treated with 1 mg/mL RNase A (Sigma) for 30 min at RT, as previously described (Beltran et al., 2016).

### Cellular transfection

Cellular transfections were performed with Lipofectamine 2000 (Invitrogen) in Serum-free Opti-MEM (GIBCO), following manufacturer instructions.

For RNA knockdown, siRNAs or Antisense Oligonucleotides were transfected for 24 hours at final concentrations of 40 nM or 80 nM, respectively, except for the 48 hours transfection to knockdown ORC1. siRNAs were designed using the i-Score designer tool and purchased from Sigma (**Table S3**). Control ASO was designed and synthesized by Ionis Pharmaceuticals. anti-TTC and anti-GAA ASOs were self-designed and synthesized by iDT, with six 2′-o-methoxyethyl nucleotides on the 5′ and 3′ ends, and nine consecutive oligodeoxynucleotides to support RNase H activity, as previously described (Zheng et al., 2010) (**Table S3**).

For exogenous ORC1 expression, pcDNA3.1 or pBABE vectors containing codon-optimized wild type or mutated cDNA sequences of ORC1, tagged with 3xFlag or Halo (**Table S2**), were transfected for 48 hours, while, in rescue experiments, plasmid transfection was preceded by 24 hours endogenous ORC1 siRNA-mediated depletion (**Table S3**). Exogenous *CEP95* was overexpressed by 48 hours transfection of a pcDNA3.1 plasmid purchased from GenScript (**Table S2**).

### Cellular synchronization and cell cycle analysis

For cellular synchronization at G1/S, U2OS cells were subjected to serum starvation for 48 hours, while HCT116 cells were blocked by double thymidine shock as previously described (Chen and Deng, 2018). Synchronization of HCT116 cells was released by PBS washing and incubation in complete culture medium, after which cells were harvested at different time points (1, 2, 3, 5, 6, 7 or 9 hours) covering the entire S phase.

For cell cycle analysis, cells were fixed in PBS 2% PFA, and incubated in 2N HCl 0.5% Triton for 30 minutes. Followed by 0.1 M Na_2_B_4_O_7_ incubation, cells were treated with RNAse A (Promega), resuspended in PBS, and the DNA stained with propidium iodide 1 mg/mL (Sigma). In flow cytometer FACSCalibur, DNA staining was recorded by BD CellQuest program. Cell cycle profiles were determined by considering the amount of labelled DNA (FL2-H) per cell.

### RNA extraction, processing, and RT-qPCR

Cell preparations were fixed with TRIzol (Sigma), and RNA precipitated with isopropanol.

In bulk RNA-seq experiments of HCT116 cells treated with control or anti-GAA ASOs, total RNA extraction was followed by Turbo DNAse (Invitrogen) digestion and library preparation with TruSeq Stranded mRNA kit from Illumina. Duplicate experiments were sequenced with Illumina NextSeq 500.

For RT-qPCR, up to 1 µg RNA was treated with DNase I (Invitrogen) and reverse-transcribed using the High-Capacity cDNA Reverse Transcription Kit (Applied Biosystem) with random hexamer primers, following manufacturer instructions. The obtained cDNA was analyzed by quantitative PCR (qPCR) using iTaq Universal SYBR Green supermix (Bio-Rad) in a ViiA™ 7 Real-Time PCR System machine (Thermo-Fisher), all reactions performed in quadruplicate. *HPRT1* or *GAPDH* RNA levels were used for normalization of total and cytoplasmic cellular extracts, while *MALAT1* was used to normalize RNA levels in nuclear extracts. For RIP and UV-RIP validations, RNA levels were normalized by their levels in a 10% experimental input of nuclear RNA. Statistical differences between relative RNA levels or relative enrichments were calculated by unpaired two-tailed Student’s *t*-test. RT-qPCR primers were self-designed or designed with the Universal Probe Library Assay Design Center (Roche), and purchased from Metabion (**Table S4**).

### Protein extraction and western blot

Soluble-chromatin cellular fractionation was performed as previously described (Shoaib et al., 2018), while subcellular fractionation in cytoplasm and nucleus, and nucleoplasm if indicated, was performed as described elsewhere (Mayer and Churchman, 2017).

Proteins from cell preparations were quantified with Pierce BCA Protein Assay Kit (Thermo Fisher). Samples were run in denaturing polyacrylamide gels by electrophoresis, and then transferred to nitrocellulose membranes (Bio-Rad). Membranes were blocked with 5% milk in PBS-Tween, and incubated overnight with primary antibodies (**Table S5**). HRP-conjugated secondary antibodies (Cell Signaling Technology) on the membrane were detected with the use of enhanced chemiluminiscence (ECL) reagent (PerkinElmer) in Odyssey CLx (LI-COR), and recorded with Image Studio Lite software. Relative protein levels were obtained based on the intensity of the western blot bands using Fiji software. Quantified intensities were normalized to those of the loading reference and, if indicated, fold changes relative to other conditions were calculated. Statistical differences between normalized intensity values were calculated by paired two-tailed Student’s *t*-test.

### ORC1 RIP-seq on nuclear extracts

Native RNA immunoprecipitation was performed as previously described (Rinn et al., 2007) with minor modifications. Briefly, 40·10^6^ asynchronous HCT116 cells, untreated or transiently expressing WT-ORC1-3xFlag, were harvested and lysed. Nuclear lysates were dounced and sonicated (Bioruptor diagenode) for 10 cycles, pre-cleared with protein A/G Dynabeads, and incubated with 5 µg of control IgG (sc-2025) or antibody of interest (anti-FLAG [M2, F1804, Sigma] or anti-ORC1 78-1-172 [Bruce Stillman laboratory (Kara et al., 2015)]). Protein A/G Dynabeads were added to sequester the antibody, and washed five times. For protein analysis, 10% of beads and inputs were resuspended in 2X Laemmli sample loading buffer, and run in acrylamide gels for western blot. RNA from beads and inputs was obtained following the RNA extraction protocol, to perform RT-qPCR (methods section *RNA extraction, processing, and RT-qPCR)* or library preparation. For sequencing, samples were first treated with Turbo DNAse (Invitrogen), and libraries were prepared with the TruSeq Stranded Total RNA kit from Illumina. Sequencing of triplicate (anti-ORC1) or duplicate (anti-Flag) experiments was done with Illumina NextSeq 500.

### ORC1 IP and phosphoproteomics

Transfected WT-ORC1-3xFlag and MUT-ORC1-3xFlag proteins in HCT116 cells (**Table S2**), in control or anti-GAA knockdown conditions (**Table S3**), were immunoprecipitated from nuclear extracts of 160·10^6^ cells, as previously described (Kara et al., 2015), with minor modifications. In brief, after cellular lysis (260 mM Sucrose, 8 mM Tris-HCl pH 7.4, 4 mM MgCl_2_, 0.8% Triton X-100), nuclei were resuspended and incubated for 30 minutes in high salt buffer (20 mM Tris-HCl pH 7.5, 400 mM NaCl, 0.4% Igepal, 5 mM MgCl_2_, 0.1 mM EDTA, 1 mM CaCl_2_, 10% Glycerol, 0.1 mM DTT), supplemented with phosphatase and protease inhibitors 1x (Roche), as well as benzonase to digest DNA. After sonication (Bioruptor diagenode) for 30 cycles, NaCl and Igepal concentrations were brought down to 200 mM and 0.2%, respectively, by adding an equal volume of dilution buffer. Clear nuclear lysates were then pre-cleared with protein G Dynabeads, and incubated with 5 µg of anti-FLAG antibody (M2, F1804, Sigma), untransfected cells being the negative control. Protein G Dynabeads were added to sequester the antibody, and washed five times with complete washing buffer (20 mM Tris, 100 mM NaCl, 0.15% Igepal, 5 mM MgCl_2_, 0.1 mM EDTA, 10% Glycerol). Beads and inputs were resuspended in 2X Laemmli sample loading buffer, and run in acrylamide gels for Coomassie blue staining.

Bands at the expected molecular weight were cut, and containing proteins were subjected to mass spectrometry. The samples were reduced with 1 mM DTT for 30 minutes at 60°C and then alkylated with 5 mM iodoacetamide for 15 minutes in the dark at room temperature. Gel pieces were then subjected to a modified in-gel trypsin digestion procedure (Shevchenko et al., 1996). Gel pieces were washed and dehydrated with acetonitrile for 10 minutes followed by removal of acetonitrile, and then completely dried in a speed-vac. Rehydration was done with 50 mM ammonium bicarbonate solution containing 12.5 ng/µL modified sequencing-grade trypsin (Promega, Madison, WI) at 4°C. Samples were then placed in a 37°C room overnight. Peptides were later extracted by removing the ammonium bicarbonate solution, followed by one wash with a solution containing 50% acetonitrile and 1% formic acid. The extracts were dried in a speed-vac (∼1 hour). For the analysis, samples were reconstituted in 5 -10 µl of HPLC solvent A (2.5% acetonitrile, 0.1% formic acid). A nano-scale reverse-phase HPLC capillary column was created by packing 2.6 µm C18 spherical silica beads into a fused silica capillary (100 µm inner diameter x ∼30 cm length) with a flame-drawn tip (Peng and Gygi, 2001). After equilibrating the column, each sample was loaded via a Famos auto sampler (LC Packings, San Francisco CA) onto the column. A gradient was formed and peptides were eluted with increasing concentrations of solvent B (97.5% acetonitrile, 0.1% formic acid). As each peptides were eluted, they were subjected to electrospray ionization and then they entered into an LTQ Orbitrap Velos Pro ion-trap mass spectrometer (Thermo Fisher Scientific, San Jose, CA). Eluting peptides were detected, isolated, and fragmented to produce a tandem mass spectrum of specific fragment ions for each peptide. Peptide sequences (and hence protein identity) were determined by matching protein or translated nucleotide databases with the acquired fragmentation pattern by the software program, Sequest (ThermoFinnigan, San Jose, CA) (Eng et al., 1994). The modification of 79.9663 mass units to serine, threonine, and tyrosine was included in the database searches to determine phosphopeptides. Phosphorylation assignments were determined by the Ascore algorithm (Beausoleil et al., 2006). All databases include a reversed version of all the sequences and the data was filtered to between a one and two percent peptide false discovery rate. Position and amount of detected phosphopeptides is presented in **Data S4**.

### ORC1 UV-RIP and iCLIP on nuclear extracts

40·10^6^ asynchronous HCT116 cells were rinsed in cold PBS and irradiated with 150 mJ/cm^2^ in a Stratalinker 2400 at 254 nm. To isolate cellular nuclei, first steps of fractionation iCLIP protocol were performed (Brugiolo et al., 2017), followed by UV-RIP or iCLIP protocols.

For UV-RIP, fixed nuclear pellets of HCT116 cells transiently expressing Flag-tagged WT-ORC1 were resuspended in RIPA buffer, and sonicated (Bioruptor diagenode) for 15 cycles. Solubilized nuclear extracts were pre-cleared with protein G Dynabeads, and 5 µg of control IgG (sc-2025) or anti-FLAG (M2, F1804, Sigma) antibodies incubated overnight at 4°C. Protein G Dynabeads were then added to sequester the antibody, and washed six times. Immunoprecipitates and inputs were eluted, and Proteinase K (NEB) was incubated for 45 minutes at 45°C for protein digestion, followed by RNA extraction and RT-qPCR (see methods section *RNA extraction, processing, and RT-qPCR)*.

iCLIP data of ORC1 was generated by following the iCLIP method described elsewhere (Blazquez et al., 2018), using 10^6^ HCT116 UV-fixed nuclei previously transfected with ORC1-WT-3xFlag (n=5), ORC1-MUT-3xFlag (n=2) or untransfected cells as control (n=1) (**Table S2**). First, nuclei were lysed in 1 mL of lysis buffer (100 mM NaCl, 50 mM Tris-HCl pH 7.4, 1% Igepal CA-630, 0.1% SDS and 0.5% Na-Deoxycholate) supplemented with protease inhibitors 1x (Roche). Nuclear lysates were sonicated (Bioruptor diagenode) for 10 cycles at low intensity. Afterwards, RNase digestion was performed for 3 minutes at 37°C with 0.4 U (n=3) or 1 U (n=2) of RNase (Thermo Scientific, EN0602) in the presence of 4U of Turbo DNase (Invitrogen), to avoid DNA contamination. Cleared supernatant was immunoprecipitated overnight at 4°C with 5 µg of anti-Flag antibody (M2, F1804, Sigma). Protein-RNA complexes were visualized using pre-adenylated, infrared dye labeled L3 adaptor with the following sequence: /5rApp/AG ATC GGA AGA GCG GTT CAG AAA AAA AAA AAA /iAzideN/AA AAA AAA AAA A/3Bio/. Reverse transcription was performed using RNA-dependent retrotranscriptase Superscript IV (Invitrogen) and barcoded primers (XXXXX) containing UMIs (NNNN): /5Phos/ WWW XXXXX NNNN AGATCGGAAGAGCGTCGTGAT /iSp18/ GGATCC /iSp18/ TACTGAACCGC. Purification of cDNAs following reverse transcription and circularization was performed using AMPure XP beads (Beckman-Coulter, USA) and isopropanol. Libraries were sequenced as single end 100 bp reads on Illumina HiSeq 4000.

### Chromatin immunoprecipitation (ChIP)

40·10^6^ asynchronous HCT116 cells, stably expressing Flag-tagged wild type and mutated ORC1 (RNA-binding mutant) (**Table S2**), or transfected to knockdown or overexpress *CEP95* mRNA (**Tables S2 and S3**) and synchronized in S phase (see *cellular synchronization* section), were fixed for 30 minutes with 2 mM DSG, and then crosslinked with 1% formaldehyde for 10 minutes. Pelleted cells were lysed (5 mM Tris-HCl pH 8, 85 mM KCl, 0.5% Igepal), nuclei resuspended in RIPA buffer, supplemented with protease inhibitors 1x, and sonicated (Bioruptor diagenode) for 30 cycles. Solubilized nuclear extracts were pre-cleared with protein A/G Dynabeads, and 5 µg of control IgG (sc-2025) and anti-CDC45 (11882, Cell Signalling Technology) or anti-PCNA (ab29, abcam) antibodies incubated overnight at 4°C. Protein A/G Dynabeads were then added to sequester the antibody, and washed with low salt (0.1% SDS, 1% Triton X-100, 2 mM EDTA, 20 mM Tris-HCl pH 8, 150 mM NaCl), high salt (Low Salt buffer with 500 mM NaCl) and LiCl (0.25 M LiCl, 1% Igepal, 1% Deoxycholate, 1 mM EDTA, 10 mM Tris-HCl pH 8) buffers, supplemented with protease inhibitors 1x (Roche). Immunoprecipitates and inputs were eluted, RNA digested with RNase A (Promega) for 30 minutes at 37°C, and Proteinase K (NEB) incubated for 45 minutes at 45°C for protein digestion. Samples were de-crosslinked overnight at 65°C, followed by phenol:chlorophorm DNA extraction and ethanol precipitation.

qPCR of precipitated DNA was done as cDNA samples (see *RNA processing* section), with self-designed primers at genomic DNA replication origins or control regions, having SNS-seq data in wild type HCT116 cells as reference (**Table S4**), and purchased from Metabion. ChIP-qPCR enrichment was done compared to a 10% input, and relativized to transfection controls. Statistical differences between relative enrichments were calculated by unpaired two-tailed Student’s *t*-test.

For sequencing of CDC45 ChIP samples, libraries of duplicate experiments were generated as previously described (Blecher-Gonen et al., 2013), and sequenced with Illumina NextSeq 2000.

### Nascent Strand Preparation by λ-exonuclease method and RT-qPCR analysis

Nascent strand preparation was performed as previously described (Almeida et al., 2018), in HCT116 cells, wild type or stably expressing WT-ORC1-3xFlag or MUT-ORC1-3xFlag (**Table S2**), untreated or transfected with control or specific siRNAs or ASOs for RNA knockdowns (**Table S3**). SNS-seq from wild type HCT116 cells was performed in triplicate experiments. To assess origin activity in HCT116 cells knocked down for GAA-RNAs, duplicate SNS-seq experiments were analyzed in cells transfected with a control ASO, or the anti-GAA ASO (**Table S3**). SNS-seq from HCT116 cells stably expressing WT or MUT-ORC1 were performed in duplicates.

Briefly, genomic DNA of 1-2·10^8^ exponentially growing cells was digested with Proteinase K (NEB), precipitated with ethanol, and solubilized in TE buffer pH 8 supplemented with RNase OUT (Invitrogen) for 48 hours at 4°C. Denatured DNA was ultracentrifuged in a seven-step 5% to 20% discontinuous sucrose gradient, with SW40Ti rotor at 24000 rpm for 20 hours at 20°C. 1 mL fractions were collected, and the DNA precipitated with ethanol. 10% volume of DNA in each fraction was denatured with 0.2 M NaOH and run onto a 1% alkaline agarose gel (50 mM NaOH, 1 mM EDTA). Followed by neutralization with 1x TAE, Syber Gold (ThermoFisher) staining was used to visualize the fractionation profile with a UV-Biorad camera. Fractions of interest (0.5 – 2 kb) were treated twice with PNK (Thermofisher) and λ-exonuclease (ThermoFisher) to remove short cut DNA, and keep newly synthetized DNA, which has a 5’ RNA segment. Efficiency of λ-exonuclease digestion was evaluated by incubation of 10% reaction volume with a digested control plasmid.

qPCR of purified nascent strands was done as cDNA samples (see *RNA processing* section), with self-designed primers at genomic DNA replication origins or control regions having SNS-seq data in wild type HCT116 cells as reference (**Table S4**), and purchased from Metabion. SNS-qPCR enrichment was done compared to a 10% input of genomic DNA, and statistical differences of relative enrichments, between control or RNA knocked down cells, were calculated by paired two-tailed Student’s *t*-test. The associated *p-values* were used for heatmap representations.

For nascent strand sequencing (SNS-seq), ssDNA fragments were converted to dsDNA as previously described (Almeida et al., 2018). First, short nascent strands were digested with RNase A/T1 mixture (ThermoFisher) to eliminate both mRNAs and RNA primers, and facilitate adaptors ligation for sequencing. Samples were mixed with random hexamer primer phosphate (Roche), and a reaction with the Klenow exo-polymerase (NEB) was used to extend the primers and synthetize the complementary DNA strand. Taq DNA ligase (NEB) was used to ligate the synthetized fragments, and the dsDNA was extracted and precipitated with ethanol. For library preparation, a protocol primarily designed for High-throughput chromatin immunoprecipitation (HT-ChIP) was followed (Blecher-Gonen et al., 2013). Duplicate or triplicate experiments were sequenced with Illumina NextSeq 2000.

### RNA-FISH

Cultured HCT116 cells on coverslips, and transfected with control or anti-GAA ASOs (**Table S3**), were fixed in PBS 3% PFA, and washed in 2x SSC 50% formamide.

Meanwhile, anti-GAA FISH probes, which were self-designed and purchased from iDT, were denaturized at 92°C for 4 minutes, cooled down, and diluted in hybridization buffer (50% Deionized Formamide, 2x SSC, 10% Dextran Sulfate) to 25 nM final concentration. FISH probes were incubated on the cells overnight at 37°C in humidity.

/56-

FAM/*T*T*C*T*T*C*T*T*C*T*T*C*T*T*C*T*T*C**T*T*C**T*T*C**T*T*C**T*T*C**T*T*C**T*T*C**T*T*C** T*T*C**T*T*C*/36-FAM/

Cells were washed with 2x SCC 50% Formamide at 55°C, followed by a wash with 2x SCC at 55°C, and a wash with 2x SCC. Preparations were blocked (PBS 0.5% Tween, 10% Heat-inactivated Goat Serum, 0.5% Blocking Reagent [Roche]), and incubated with α-FAM-POD antibody (Roche). After washing with 4x SSC, preparations were incubated with TSA-Cy3 (Perkin Elmer) diluted in Amplification Diluent (Perkin Elmer). Secondary antibodies were washed with 4x SSC, followed by a wash with 4x SSC 0.1% Triton, and a last wash with 4x SSC. Cells were mounted on microscope glass slides with mounting solution with DAPI (Vectashield), and imaged with confocal fluorescence microscope Zeiss LSM 880 NLO 63x objective, and images were captured with the ZEN microscopy software (Zeiss). Fiji software was employed for stacks deconvolution and signal quantifications. Probe fluorescence of entire images (15 cells each) were compared between control and anti-GAA treated cells by unpaired two-tailed Student’s *t*-test.

### Protein imaging (chromatin immunofluorescence, STORM)

Protein imaging at chromatin of HCT116 cells, after removing soluble cellular fractions, was performed as previously described (Scovassi and Prosperi, 2006), with anti-CDC45 (Cell Signalling 11881) and anti-PCNA (sc-56) antibodies. After immunofluorescence staining with secondary antibodies, coverslips were mounted on microscope glass slides with mounting solution with DAPI (Vectashield). Preparations were imaged using the 63x objective of the automated microscope Zeiss Axio Imager M1, and images were captured with the ZEN microscopy software (Zeiss). Fiji software was employed for stacks deconvolution and signal quantifications. After delimiting cellular nuclei, fluorescence intensities were measured and normalized by nuclei area. Comparisons between cells (>100) in different experimental conditions were done by unpaired two-tailed Mann Whitney *t*-test.

In STORM experiments, synchronized U2OS cells (see *cellular synchronization* section), untreated, mock-transfected, or transiently expressing Halo-tagged wild type and mutated ORC1 (**Table S2**), were permeabilized before fixation, to capture protein and RNA imaging at chromatin, as previously described (Chen et al., 2015; Scovassi and Prosperi, 2006). Incorporated EU (see *culture treatments* section) was detected with the Click-it Plus Kit (ThermoFisher) AF-647 (ThermoFisher), and ORC1 with anti- ORC1 antibody (F-10) (sc-398734) or Janelia Fluor® 549 HaloTag® Ligand (Promega, GA1110). After immunofluorescence staining with secondary antibodies, coverslips were mounted on microscope glass slides with freshly prepared super resolution imaging buffer (PBS 1 mg/mL Glucose Oxidase [Sigma], 0.02 mg/mL Catalase [Sigma], 10% Glucose [Sigma], 100 mM Mercaptoethylamine [Thermo Fisher]) flowed through. All raw images were acquired using a custom-built inverse microscope platform (Applied Scientific Instrumentation). Briefly, 639 nm (UltraLaser, MRL-FN-639-1000) and 561 nm (Coherent, Sapphire 561 LPX - 500) laser lines were adjusted to 1.5 and 0.8 kW/cm^2^, respectively. Fluorescence emission was expanded with a 1.67× achromatic lens tube and was collected on a sCMOS camera (Photometrics, Prime 95B). Fluorescence signals were collected sequentially using the AF647 (Semrock, FF01-676/37) and AF568 (Semrock, FF01-607/36) single-band pass filters in a filter wheel (ThorLabs, FW102C). A 405 nm laser line (UltraLaser, MDL-III-405-500) was introduced to enhance recovery of dark state fluorophores when required. 2000 Frames at 33 Hz were acquired for each color.

Localization of single molecules and the mapping of the two different channels were carried out in algorithms written in MATLAB as previously described (Yin et al., 2019). To quantify the degree of colocalization between RNA and ORC1, cross-correlation (Equation (1)) between the two species was calculated. Similar to the radial distribution function, for two images *Im_1_* and *Im_2_*, the correlation magnitude at displacement **r** = (*r, θ*) is defined as

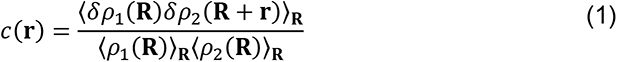

where *ρ_i_* (**R**) denotes the local density of *Im_1_* at location **R** and 〈*ρ_i_*(R)〉_R_ denotes the average density over the entire image, where <·>_R_ denotes the average operator over all the location **R**; *δρ_i_*(R) = *ρ_i_*(R) – 〈*ρ_i_*(R)〉_R_ denotes the fluctuation of the local density at location **R**. As the correlation is not orientation-specific, the 2D *c*(**r**) = *c*(*r*,*θ*) was further averaged over *θ* and plotted as the correlation profile as the function of the radial distance *r*.

In brief, each nucleus was first manually outlined to generate a ROI for independent analysis. EU and ORC1 signals from the same ROI were submitted for cross-correlation analysis to obtain their association magnitudes, whilst the cross-correlation between the two species from different ROIs serve as a control describing random distributions (Chen et al., 2015). Unpaired two-sample *t*-test between experimental and randomized data was done to determine the significance of the correlation. Same analyses on cells with no EU incubation were used as the experimental negative control.

### Fiber stretching and staining

500 HCT116 cells transfected with siRNAs (**Table S3**), plasmids to express exogenous ORC1 (**Table S2**), or ASOs (**Table S3**), and pulsed with CldU and IdU (see *culture treatments* section), were dropped on Superforst Thermo Scientific microscope slides and lysed with spreading buffer (0.5% SDS, 200 mM Tris-HCl pH 7.5, 50 mM EDTA) in humidity. Slides were tilt at a 10-15° angle to allow the DNA suspension to run slowly down the slide, and air dried. DNA fibers were fixed in -20° cold 3:1 methanol:acetic acid, and air dried. For fiber staining, DNA was denatured in 2.5 M HCl for 30 minutes at room temperature, washed with PBS, and blocked (PBS 1% BSA, 0.1% Triton). Slides were then incubated with primary antibodies detecting CldU (ab6326, abcam) and IdU (347580, BD), in a humidity chamber overnight at 4°C. After PBS washing, slides were incubated with fluorescent secondary antibodies. After washing, primary anti-ssDNA (MAB3034) and secondary fluorescent antibodies to label DNA fibers were incubated for 30 minutes. Preparations were then air dried and mounted with Prolong diamond (Invitrogen). Preparations were imaged using the 40x objective of the automated microscope Zeiss Axio Imager M1, and images were captured with the ZEN microscopy software (Zeiss).

DNA fiber images were analyzed with Fiji software, considering a conversion factor of 1 µm = 2.59 kb. Two parameters were analyzed: fork rate, measuring the length (in kb) of the IdU track and dividing it by the 20 minutes of the duration of the pulse; inter origin distance, measuring the distance between adjacent origins (recognized as IdU-CldU-IdU tracks). Comparisons of fork rate and inter origin distances between experimental conditions were analyzed by unpaired two-tailed Mann Whitney *t*-test.

### Protein purification

Purified wild type and mutated (R441A, R444A, R465A) ORC1 RNA-binding regions (amino acids 413-511), fused to GST, were produced in Shou Waga laboratory (Hoshina et al., 2013). Protein concentrations were estimated by Coomassie blue or silver staining, compared to BSA known concentrations.

Wild type and mutated full length MBP-PP-GFP-ORC1-6xHis were expressed and submitted to Ni-NTA beads purification followed by a second purification with amylose beads, as previously described (Hossain et al., 2021). Briefly, *E.coli* BL21 cells were transformed with fusion plasmids (tagged full-length ORC1) (**Table S1**), grown in LB media at 37°C until 0.7-0.9 O.D., and induced overnight with 0.3 mM IPTG at 16°C. Bacterial cells were then pelleted, washed, and lysed with 100 mg/mL lysozyme in buffer A (25 mM Tris pH 7.5, 150 mM NaCl, 0.02% Igepal, 5 mM MgCl_2,_ 5 mM Benzamidine-HCl, 1mM PMSF, 1x protease inhibitors [Roche], 10% Glycerol). After centrifugation, the clarified supernatant was incubated with pre-washed Ni-NTA beads for 3 hours at 4°C. Bead bound proteins were washed with lysis buffer, and eluted with 300 mM imidazole. WT and MUT MBP-PP-GFP-ORC1-6xHis proteins were further purified with amylose beads and eluted with 20 mM maltose. Protein concentrations were estimated by Coomassie blue or silver staining, compared to BSA known concentrations.

### In vitro assays (EMSA, LLPS, in vitro phosphorylation, binding assays)

In electrophoretic mobility shift assays (EMSA), wild type and mutated GST-ORC1 (413-511) (concentrations indicated in figure captions) or control buffer (25 mM HEPES pH 8, 300 mM NaCl, 10% Glycerol, 1 mM DTT), were incubated with RNA (concentration indicated in figure captions) in 20 µL binding buffer (25 mM HEPES pH 8, 10 mM Mg(C_2_H_3_O_2_)_2_, 0.1 mM EDTA, 5% Glycerol), supplemented with BSA 2 mg/mL, 3 mM ATP (NEB) and RNAsin ribonuclease inhibitors (Promega). Binding reaction was incubated at 30°C for 30 minutes, and samples were immediately loaded into a pre-run non-denaturing gel for electrophoresis at 4°C. RNA was stained in Sybr Gold (ThermoFisher), visualized with a UV-Biorad camera, and quantified with Fiji software. In experiments with total RNA, RNA fragments were obtained by sonicating (Bioruptor diagenode) total extracts of HCT116 cells, and purifying short RNA fragments with miRNA columns (PureLink). Electrophoresis was done in 7.5% polyacrylamide gels. In EMSAs with *CEP95* RNA, RNA fragments were *in vitro* synthetized with T7 RNA polymerase (Promega), using as a template the PCR products amplified from the pCDNA3-CEP95 vector (**Tables S2 and S4**). Electrophoresis was done in 0.7% agarose gels.

In liquid-liquid phase separation (LLPS) assays, full length wild type and mutated MBP-PP-GFP-ORC1-6xHis (4 µM) were resuspended in LLPS reaction buffer (50 mM Tris pH 7.5, 100 mM NaCl, 1 mM MgCl2, 1 mM DTT) supplemented with Prescission protease, in presence or absence of 4 µM of RNA probes previously used (same sequence) in DNA phase separation experiments (Hossain et al., 2021; Parker et al., 2019) (GAAGCTAGACTTAGGTGTCATATTGAACCTACTATGCCGAACTAGTTACGAGCTAT AAAC), and incubated at 4°C for 16 hours. Following incubation, the reactions were spotted on microscope slides with coverslips and observed immediately with 63x oil immersion objective of the automated microscope Zeiss Axio Imager M1. Images were captured with the ZEN microscopy software (Zeiss). GFP-droplet size was measured with Fiji software.

### CatRAPID analysis

The *cat*RAPID algorithm estimates the binding potential of a protein-RNA pair through van der Waals, hydrogen bonding and secondary structure propensities, allowing identification of binding partners with accuracy of 0.78 or higher (Bellucci et al., 2011; Cirillo et al., 2016).

The *cat*RAPID analysis to predict ORC1 direct interactions with ORC1 RIP-RNAs was performed following standard pipelines (Agostini et al., 2013). Briefly, we used the major RNA isoform for each gene reported in ORC1 RIP-seq experiments, retrieved the ORC1 interaction scores from RNAct (Lang et al., 2019), and all transcripts with *p*-*value* > 0.01 were filtered out. Two classes were analyzed: depleted RNA, when *log2 Fold Change* < 0, and enriched RNA, if *log2 Fold Change* > 0, and the difference in the predicted *cat*RAPID scores of enriched and depleted was represented (z-score) and computed with unpaired two-tailed Student’s *t-*test. We also used the experimental *log2 Fold Change* to rank the two groups of transcripts, enriched and depleted. Equal fractions of enriched (i.e., highest fold changes) and depleted (i.e., lowest fold changes) RNAs were compared. The discriminative power, measured as the Area under the ROC curve (AUC), increases proportionally to the experimental signal and reaches a value of 0.80, indicating strong enrichment of predicted physical interactions.

*cat*RAPID *omics v2* was used to predict RNA interactions of the wild type and mutated ORC1 protein sequences (R441A, R444A and R465A) (Armaos et al., 2021). We divided the Interaction Propensity score in bins of width 20 (a.u.). For each of the Interaction Propensity Bins, we calculated the fraction of RNAs that obtained decreased Interaction Propensity score against the mutated ORC1 sequence. We found that the RNAs exhibiting ‘High Interaction Propensity’ scores were enriched targets of WT- ORC1, while those exhibiting ‘Low Interaction Propensity’ scores were enriched targets of MUT-ORC1. This finding reinforces the notion that the identified RNAs by RIP-seq are bona-fide ORC1 direct interactors.

### cleverSuite analysis

To investigate the effect of amino acid mutations on ORC1, we used the *CleverSuite* approach (Klus et al., 2014), which uses two protein sequence sets (Positive and Control/Negative) to build a model able to separate them based on physicochemical features (hydrophobicity, secondary structure, charge, etc). The model can be reused to predict classification of other sets. We trained 2 models: RNA and DNA binding ability. For the RNA model we used RNA binding proteins (Castello et al., 2012) and, as control, a set of proteins that were found in the lysate from the same study. For the DNA model we used Proteins annotated as “DNA binding” from UniProt and, as control, a sample set of similar size as the DNA binding proteins that were not annotated as “DNA binding” nor “RNA binding”. The two models can be found at the links:

http://crg-webservice.s3.amazonaws.com/submissions/2021-09/393018/output/index.html?unlock=8bc230ac56

http://crg-webservice.s3.amazonaws.com/submissions/2013-12/17868/output/index.html?unlock=f3a7ffa08f

We used the predictive ability of the *CleverSuite* to assess if the ORC1 mutant (R441A, R444A and R465A) belongs to the positive or control set. Specifically, we tested the hypothesis of whether ORC1 region including the mutations affected its ability to interact with RNA (positive) and/or DNA (control). The results unequivocally showed that mutant ORC1 has decreased RNA binding activity, while its DNA binding activity remains unaffected. In the analysis we considered different ORC1 fragments centered around the mutations to control the signal to noise ratio of *CleverSuite* scores. Specifically, in the RNA Binding model, all ORC1 WT fragments are predicted as “Positive” (RNA binding) and all the Mutated ORC1 fragments were predicted as “Control”. Importantly, the same regions were predicted all as “Positive” for both WT and mutant (DNA binding) when using the second (Control) model. Thus, the DNA binding ability is predicted to be unaffected by the RNA-binding mutations.

### Statistical analysis

Experimental data was plotted and analyzed using the GraphPad statistical software, following the statistical analysis for each type of data, specified in each method section and/or figure captions. Most experimental data is represented as the average of at least three biological replicates, indicated at figure captions. Imaging data is presented as a representative experiment with multiple measurements, which was validated in additional biological replicates. Number of replicates in sequencing experiments is specified in each method section.

R software was used for bioinformatic analysis, using R package ggplot2 to generate different types of plots (https://cran.r-project.org/web/packages/ggplot2/index.html). Significance was obtained using the statistical test corresponding to each type of data analyzed as explained in each analysis section.

In all cases, *p-values* were given using the following thresholds: ns for *p-value* > 0.05; * for *p-value* ≤ 0.05; ** for *p-value* ≤ 0.01; *** for *p-value* ≤ 0.001.

### RNA-seq, RIP-seq and ChIP-seq Pipelines

In ASO control and anti-GAA (**Table S3**) RNA-seq of HCT116 cells, and ORC1 and ORC1-3xFlag RIP-seq experiments, QC of sequencing files was performed with FastQC (http://www.bioinformatics.babraham.ac.uk/projects/fastqc/). Fastq files were aligned with STAR (Dobin et al., 2013), reads aligning to GL contigs were removed, and FeatureCounts (Liao et al., 2014) was used to quantify the number of reads falling in annotated genes in hg19 human reference genome, downloaded from Ensembl (Yates et al., 2020). DESeq2 (Love et al., 2014) was used to measure differential expression, being ASO control and inputs the reference conditions for RNA-seq and RIP-seq analyses, respectively.

RNA-seq public sequences from untreated HCT116 cells (Gajdušková et al., 2020) (GSE118051) were aligned to the hg38 reference genome, downloaded from GENCODE (Frankish et al., 2019), using STAR (Dobin et al., 2013) with parameters: ‘winAnchorMultimapNmax20 -outFilterMultimapNmax 20 -twopassMode Basic’. Public ChIP-seq sequencing reads from duplicate experiments (1,2) were analyzed using a Nextflow pipeline (DI Tommaso et al., 2017) with golden standard of the NF-Core consortia (https://nf-co.re/). From ENCODE (Consortium, 2004): H3K27me3 (ENCFF457PEW), H3K9me3 (ENCFF020CHJ), H3K4me1 (ENCFF531IUP), H3K27ac (ENCFF227RRY), H3K4me3 (ENCFF213WKK), H3K36me3 (ENCFF059WYR). From public data(Long et al., 2020): H2A.Z (SRR9850576, SRR9850577), H4K20me1 (SRR9850580, SRR9850581), H4K20me2 (SRR9850582, SRR9850583), ORC1 ChIP input (SRR9850586) and IP (SRR9850594, SRR9850595). Raw sequencing reads were trimmed and quality control was performed to remove poor quality sequences. Adapter-trimmed reads were then aligned to hg38 human reference genome, downloaded from GENCODE (Frankish et al., 2019), with BWA (Li and Durbin, 2010) under default parameters. Replicates were merged and deduplicated to remove optical reads ([CSL STYLE ERROR: reference with no printed form.]). To obtain robust estimates of the results, we respected the best practices as suggested by ENCODE consortia (Landt et al., 2012). Then, aligned reads passing all ChIP-seq QC metrics were submitted to MACS2 peak calling (Zhang et al., 2008) comparing no antibody input and corresponding samples using the function ‘callpeak -g hs -B -q 0.05 --fe-cutoff 1.5 – broad’.

Self-generated CDC45-treated ChIP-seq samples were trimmed to remove adapted and low-quality sequenced reads. Bowtie2 (Langmead and Salzberg, 2012) was used to align the reads to the hg19 human reference genome downloaded from ENSEMBL (Yates et al., 2020). Coverage tracks were generated with bamCoverage (Ramírez et al., 2014), and the average signal at TSS positions (-/+ 5kb) was divided in bins of 10bp length, with computeMatrix from deeptools, to be analyzed by paired *t*-test (Euclidean distance of 14.0163511182744 and 13.8052506781907 for replicates 1 and 2, respectively; *p-value* 3.08e-16 and 7.73e-65 for replicates 1 and 2, respectively). Coverage differences (and associated *p-values*) between WT and MUT-ORC1 cells were also measured at TSSs of genes in individual iCLIP-defined quantiles.

### iCLIP analysis

ORC1 iCLIP sequencing reads were analyzed on the iMaps server (<u>Genialis Workspace</u>) using the iCount software (Ule et al., 2010) (https://github.com/tomazc/iCount). Briefly, experimental barcodes were removed and sequencing reads aligned with STAR (Dobin et al., 2013) to hg38 human reference genome downloaded from GENCODE (Frankish et al., 2019), allowing two mismatches and ten secondary alignments. DNA or chromatin contamination was excluded by aligned data interrogation with infer_experiment.py package from RSeQC software (Wang et al., 2012). Unique Molecular Identifiers (UMIs), were used to distinguish and remove PCR duplicates. To determine protein-RNA contact sites, the sequencing read preceding nucleotide was allocated as the crosslink site event.

The presented data refers to analysis from the three experimental replicates of ORC1- 3xFlag transfected cells and low (0.4 U) RNase treatment, while non transfected control was used to corroborate data specificity (**Table S1**). Replicates were merged and summary of cDNA counts within genes and genic regions were generated with iCount summary function. Assignment of crosslink sites to coding transcripts, non-coding or biotype features, was done by following segmentation hierarchy rules (https://github.com/tomazc/iCount/blob/master/iCount/genomes/segment.py).

For data representation, iCLIP signal was normalized by sequencing deep and million of tags (CPM) and binned per nucleotide. Coverage tracks were generated using deepTools (Ramírez et al., 2014), and metagene plots were drawn using normalized coverages between the transcriptional start site (TSS) and the transcriptional termination site (TTS) of genes, defined by the GENCODE (Frankish et al., 2019) annotation from hg38 human reference genome, using 100 nucleotide bins. Normalized iCLIP data was also plotted in metagenes, in -/+ 10kb windows around genomic TSS positions.

The significant crosslink signal was normalized by sequencing deep and million of tags (CPM). Significant contact sites were identified as iCLIP peaks, using the iCount peak function, based on false discovery rate (FDR) <0.05 comparing specific sites within a window of three nucleotides with randomized data (100 permutations) and within co-transcribed regions (https://github.com/tomazc/iCount/blob/master/iCount/analysis/peaks.py).

To identify RNA motifs mediating ORC1 binding, iCLIP peaks with more than 5 crosslinks per nucleotide were slop 100 nucleotides both sides and submitted to MEME motif finding algorithm (Bailey and Elkan, 1994) with parameters ‘-rna -maxw 6 -maxsize 1000000000 -neg ’, comparing positive sites with negative randomized data from SNS-seq experiments and from homologous genomic regions. Although several motif sizes were tested, 6-mers appeared to be the most reliable. G4 predictions were also obtained, by using TetraplexFinder from the QuadBase2 web server (quadbase.igib.res.in) (Dhapola and Chowdhury, 2016), using pre-set motif configuration (medium stringency G3 L1-7, Greedy search algorithm, Bulge size=0) and search ’+’ strand only. Statistical significance was analyzed by two-proportions z-test. ORC1 binding preferences for small nuclear RNAs (snoRNAs) annotated hg38 reference genome, downloaded from GENCODE(Frankish et al., 2019), were separately and further assessed in experimental snoDB data base v.1.2.1 (http://scottgroup.med.usherbrooke.ca/snoDB/ downloaded in December 2020).

### ORC1 interactome analysis (iCLIP-RIP comparison, genomic data, MEME motifs)

SAMtools (Li et al., 2009) and Bedops (Neph et al., 2012) were used to do different types of operations with genomic data, and the UCSC liftOver tool was used to convert coordinates between different genome versions.

Effective ORC1 protein RNA contact sites from iCLIP were pooled overlapping Ensembl IDs from iCLIP peaks (> 5 crosslink sites and < 0.05 FDR) and RIP-seq technique (*log2 Fold Change* > 1 and *p-value* < 0.05) based on annotation of hg38 human reference genome, downloaded from GENCODE (Frankish et al., 2019), defining ORC1 iCLIP- RNAs, RIP-RNAs, ORC1-RNAs (union iCLIP and RIP) or HC ORC1-RNAs (overlap iCLIP and RIP) (**Data S2**). Hypergeometric test confirmed the significance of the overlap and the union of both techniques, also when applying different iCLIP cut-offs. Furthermore, to study general agreement between ORC1 RIP and iCLIP in identifying the same groups of transcripts, ORC1 bound and not bound groups of transcripts were defined by iCLIP CTPM parting from general transcriptomic data in HCT116 cells (Gajdušková et al., 2020). Statistical differences between these two groups of transcripts in terms of CTPM and RIP enrichment (*log2 Fold Change*) were analyzed by unpaired two-tailed Student’s *t*-test.

Biotypes and gene length of different groups of genes (ORC1-RNAs, HC ORC1-RNAs, RIP-RNAs, iCLIP-RNAs) were obtained from Ensembl BioMart (Yates et al., 2020), and HCT116 expression data were obtained from Array Express (Athar et al., 2019) study E-MTAB-2770 (RNA-seq of 934 human cancer cell lines from the Broad-Novartis Cancer Cell Line Encyclopedia (Barretina et al., 2012)). Negative controls of ORC1-RNAs and HC-ORC1-RNAs consisted in an equal number of RNAs with *log2 Fold Change* < -0.25 and *p-adj* < 0.5, or with -0.16 < *log2 Fold Change* < 0.16 in ORC1 RIP-seq experiment, to consider genes with no ORC1 RNA-binding, with expression in HCT116 cells. Comparisons of gene length and expression between the different groups of genes were statistically interrogated by unpaired two-tailed Student’s *t*-test. Same controls were used to determine whether ORC1-RNAs and HC ORC1-RNAs interact with each other in the 3D nucleus with different frequencies than controls, and Hi-C data of HCT116 cells were obtained from GEO series GSE104333 (Rao et al., 2017) (untreated synchronized combined MAPQ >= 30). Juicer (Durand et al., 2016) tools command dump was used to extract data from the .hic file and obtain KR normalized intra-chromosomal contact information at 100 kb resolution. Statistical significance in terms of Hi-C contacts between control or genes of interest was analyzed by two-proportions z-test.

To establish ORC1-RNAs localization within early or late replicating regions of the genome, Repli-seq data for HCT116 cells were obtained from ReplicationDomain (https://www2.replicationdomain.com/index.php) (Weddington et al., 2008) database, hg19 human reference genome (Int90617792, Int97243322). Significant enrichment of ORC1-RNA and HC ORC1-RNA genes for early-replicating regions was assessed by hypergeometric test, considering the universe of early and late replicating regions genome-wide.

To detect enriched sequence motifs through entire ORC1-bound RNAs, sequences of the largest transcripts were extracted from the list of candidate genes (**Data S2**). Input datasets of RNA sequences were run to find significantly enriched motifs on any length with MEME (Bailey and Elkan, 1994), with respect to a control dataset of the same number of transcript sequences (option -neg) by using the differential enrichment objective function. Negative controls consisted in the same number of randomly selected transcripts from ORC1 RIP-seq data, thus expressed in HCT116 cells.

### SNS-seq analysis

The quality of the sequencing files was assessed with FastQC (http://www.bioinformatics.babraham.ac.uk/projects/fastqc/). Bowtie2 (Langmead and Salzberg, 2012) was used to align the reads to the human hg19 reference genome, downloaded from Ensembl (Yates et al., 2020), and Picard (http://broadinstitute.github.io/picard) was used to remove duplicate reads.

To compare SNS-seq in control vs anti-GAA cells, statistical analyses were done for each of the experimental replicates, by considering the normalized raw sequencing signal (bamCoverage, RPKM normalization) at TSS (-/+ 7.5 kb) genomic positions. The average signal was divided in bins of 10bp length, with computeMatrix from deeptools, and analyzed by paired *t*-test (Euclidean distances of 1.471843 and 2.139965, for replicates 1 and 2, respectively; Fold change ASO anti-GAA vs ASO control of 0.9872622 and 0.9704698, for replicates 1 and 2, respectively; *p-value* 8.85552e-36 and 6.840894e-167 for replicates 1 and 2, respectively).

To compare SNS-seq experiments in WT vs MUT-ORC1 expressing HCT116 cells, we merged raw sequencing signal (bamCoverage, RPKM normalization) at TSS (-/+ 5 kb) genomic positions, from the two experimental replicates. The average signal was divided in bins of 10bp length, with computeMatrix from deeptools, and compared by paired *t*-test (Euclidean distance of 2.96711709492902; Fold change WT vs MUT 1.059799; *p-value* 2.2e-16).

Origin peaks were determined following the analysis pipeline in (Langmead and Salzberg, 2012), that uses two peak callers: MACS2 (Zhang et al., 2008) with a threshold of q=0.1 to identify narrow peaks, and EPIC2 (Stovner and Sætrom, 2019) with FDR=0.1, to detect diffuse peaks. To obtain the common peaks detected with both methods, we used intersectBed from BedTools (Quinlan and Hall, 2010), with parameters -wa -u (nonreciprocal, report any overlapping features). The final set of peaks considered for subsequent analyses includes those common MACS2+EPIC2 peaks that are present in at least two out of the three SNS-seq replicates in the wild type HCT116 samples, and in both replicates anti-GAA and control ASOs.

SNS-seq identified 37725 origins that were consistent with previously published origin mapping in other cell types (Akerman et al., 2020), since 63% of the HCT116 origins overlapped with those in quantiles Q1 and Q2 of the 10 defined by the mentioned study, which represent the most robust origins with the highest conservation among cell types (called core origins) (Akerman et al., 2020). Significant enrichment of SNS-seq peaks at TSSs of ORC1-RNA genes, also visualized in metagenes of normalized reads, was assessed by hypergeometric test, considering the presence of SNS-seq peaks at TSS of genes in the entire genome. To compare anti-GAA samples against control samples, peaks were divided in different groups: present only in control replicates, present only in anti-GAA replicates, common peaks (those overlapping anti-GAA and control peaks), and peaks differentially bound in anti-GAA or in control. Differentially bound peaks were determined with DiffBind (Stark and Brown) (v.2.10.0) and DESeq2 (Love et al., 2014) (v.1.22.1) R packages. ChiPseeker (Yu et al., 2015) was used for SNS-seq peak annotation, comparison and visualization in heatmaps around TSS positions (13108 peaks in ASO control; 12198 peaks in ASO anti-GAA).

Differences in SNS-seq (ASO control vs ASO anti-GAA, and WT-ORC1 vs MUT-ORC1) in combination with other data (RNA-seq ASO control vs ASO anti-GAA, or ORC1 iCLIP) were assessed by Gene Set Enrichment analyses (GSEA, see *combined data and correlation analyses* section).

### Combined data and correlation analyses (ORC1 iCLIP, RIP-seq, SNS-seq, RNA-seq)

ChIP-seq, SNS-seq and RNA-seq data were normalized by RPKM, in non-overlapping bins of 10 nucleotides. Genome-wide tracks of normalized ORC1 iCLIP (crosslinks and iCLIP peaks), SNS-seq, ORC1 RIP-seq, and public ChIP-seq data (Long et al., 2020), were visualized and plotted in the IGV (integrative genomics viewer) browser (Robinson et al., 2011). Normalized sequencing data was also presented in coverage tracks and metagenes, using deepTools (Ramírez et al., 2014), heatmaps, and violin or bar plots.

To study the correlation between replication origins and chromatin marks at TSSs of genes encoding for ORC1-RNAs, normalized ChIP-seq, SNS-seq and RNA-seq data were represented in a correlation heatmap, with associated Spearman’s Correlation values.

ORC1 RNA-binding iCLIP data, normalized by sequencing deep and million of tags and binned per nucleotide (CTPM), was used to define the division of ORC1-RNA genes (union RIP-seq and iCLIP – see *Genomic data analysis* section) in 6 quantiles (Q1 to Q6), defining levels of direct ORC1-RNA binding. Having determined iCLIP-defined groups of genes, normalized sequencing data was represented and subjected to statistical analyses, to study the correlation between replication origins, chromatin marks, and ORC1 RNA-binding. Statistical differences between quantiles of genes in terms of RNA levels (RNA-seq), SNS-seq and ChIP-seq, represented in profile, heatmap, violon or bar plots, were analyzed by unpaired two-tailed Student’s *t*-test.

Gene Set Enrichment Analysis (GSEA) analyses were performed using fgsea (Korotkevich et al., 2021) (v.1.22.0) R package with 10,000 permutations to calculate statistical significance. Genes were filtered ranked according to their *log2 Fold Change* between anti-GAA vs Control RNA-seq samples, or between the signal across the TSS (-/+ 5Kb) in WT-ORC1 against MUT-ORC1 SNS-seq replicates. In both cases, genes with significant (adjusted *p*-*value* < 0.05) differences between conditions were considered for generating the ranking lists.

Ranked genes according to their upregulation or downregulation in RNA-seq experiments (ASO control vs ASO anti-GAA) were crossed with a list of genes with reduced SNS-seq signal in the anti-GAA condition. This second list of genes was defined by DiffBind (Stark and Brown) (v.2.10.0), with a cut-off of p-value < 0.05, to select genes with TSS SNS-seq peaks (see *SNS-seq analysis* section) enriched in the control condition.

Ranked genes according to their enrichment of SNS-seq signal at TSS (WT vs MUT SNS-seq) were crossed with the list of genes in iCLIP-defined quantiles, individually (Q1, Q2, Q3, Q4, Q5 and Q6) or in combination (Q1+Q2+Q3 and Q4+Q5+Q6).

### In silico analysis of ORC1 sequence, structure, and conservation

To trace orthologues of human ORC1, sequence datasets for 132 proteomes (**Data S4**) were downloaded from the available databases comprising 53 prokaryotes and 79 eukaryotes.

Homologous sequences of human proteins were identified using Inparanoid (Remm et al., 2001), an automatic method that uses pair-wise similarity scores between two proteomes for constructing orthology clusters, calculated using NCBI-Blast. The program was run using default parameters except for the in-paralog confidence cut-off, which we made more stringent (from 0.05 to 0.25). All Inparanoid blasts were run using a threshold *e-value* of 0.01 and different matrices were used in pair-wise comparisons to account for different evolutionary distances: Blossum45 to compare prokaryotes, Blossum62 for eukaryotes, and Blossum80 for comparisons between metazoans. The L-INS-i model in Mafft (Katoh and Standley, 2013) was used to build a multiple sequence alignment (MSA) with the ORC1 orthologous proteins from vertebrates and metazoans (**Table S5**). The alignment was visualized using Jalview (Waterhouse et al., 2009) and its quality was manually checked. Consensus sequences logos were generated with WebLogo (Crooks et al., 2004). We used MEME Suite web platform (Bailey et al., 2009) to find motifs within the consensus sequence of the vertebrate RNA-binding sequence of ORC1 orthologues and MEME FIMO(Grant et al., 2011) to search for the TPR/K motif in the *H. sapiens* genome, Ensembl (Yates et al., 2020). To analyze the domain repertoire of ORC1 orthologues, we ran the Hmmscan program from HMMER 3.2 (hmmer.org) (Eddy, 2011) against the Pfam database (version 32, September 2018) (El-Gebali et al., 2019). Non-overlapping hits with scores above the conditional e-value threshold of 0.05 were considered significant.

ORC1 protein secondary structure was predicted with PsiPred (Jones, 1999), and the MetaDisorder server (Kozlowski and Bujnicki, 2012) was used to predict intrinsic protein disorder using iPDA (Su et al., 2007), PrDOS (Ishida and Kinoshita, 2007), Pdisorder (http://www.softberry.com/) and IUPred long (Dosztányi et al., 2005). Phyre2 (Kelley et al., 2015) was used to predict the 3D protein structure of human ORC1 and the model was visualized and colored using the PyMOL Molecular Graphics System (Schrödinger, LLC.).

**Data S1 (separate file). RIP-seq data (ORC1 and ORC1-Flag).** RNAs identified by RIP-seq of endogenous and exogenous ORC1 are listed, showing fold changes and *p-values* of replicate experiments.

**Data S2 (separate file). ORC1 RIP-seq and iCLIP data. (A)** ORC1-RNAs are listed, showing gene identification and characteristics, RIP counts and enrichment, and iCLIP crosslink and peak counts. **(B)** HC-ORC1 RNAs are listed.

**Data S3 (separate file). Phosphopeptides detected by mass spectrometry.** Position of phosphorylated residues in WT and RNA-binding mutant (MUT) ORC1, in control (ASOC) or GAA-RNA (ASO anti-GAA) conditions.

**Data S4 (separate file). Sequence datasets used to identify ORC1 human orthologues.** 132 proteomes and taxonomy are listed.

---

**Figure S1.**
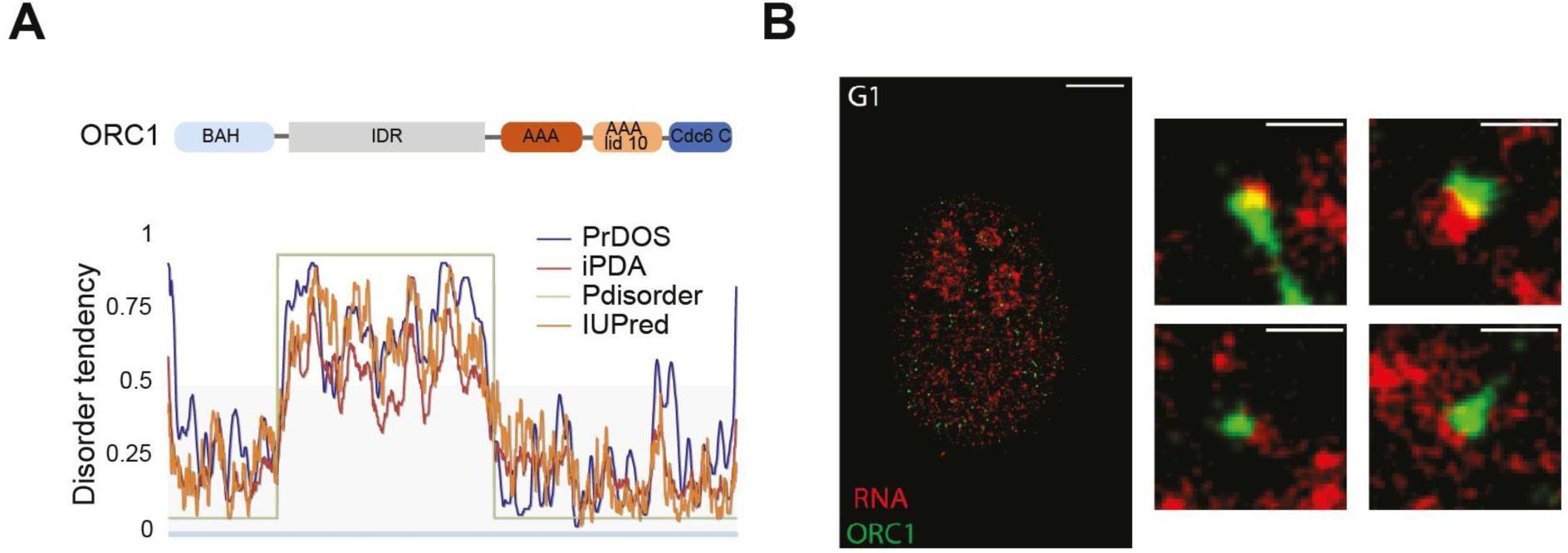
ORC1 protein structure and colocalization with RNA in cells. **(A)** Representation of linearized human ORC1 showing canonical protein domains, as defined by Pfam (El-Gebali et al., 2019), and the central IDR. Below, disorder plot showing ordered or disordered regions (Disorder Tendency < or > 0.5) along protein residues (x axis), defined by disorder predictors (colored lines). **(B)** Representative STORM images of U2OS nucleus labelled with chromatin-associated RNA (long pulse) and ORC1 (Scale bar, 2 μm), and representative zoom-in foci showing their colocalization.

**Figure S2.**
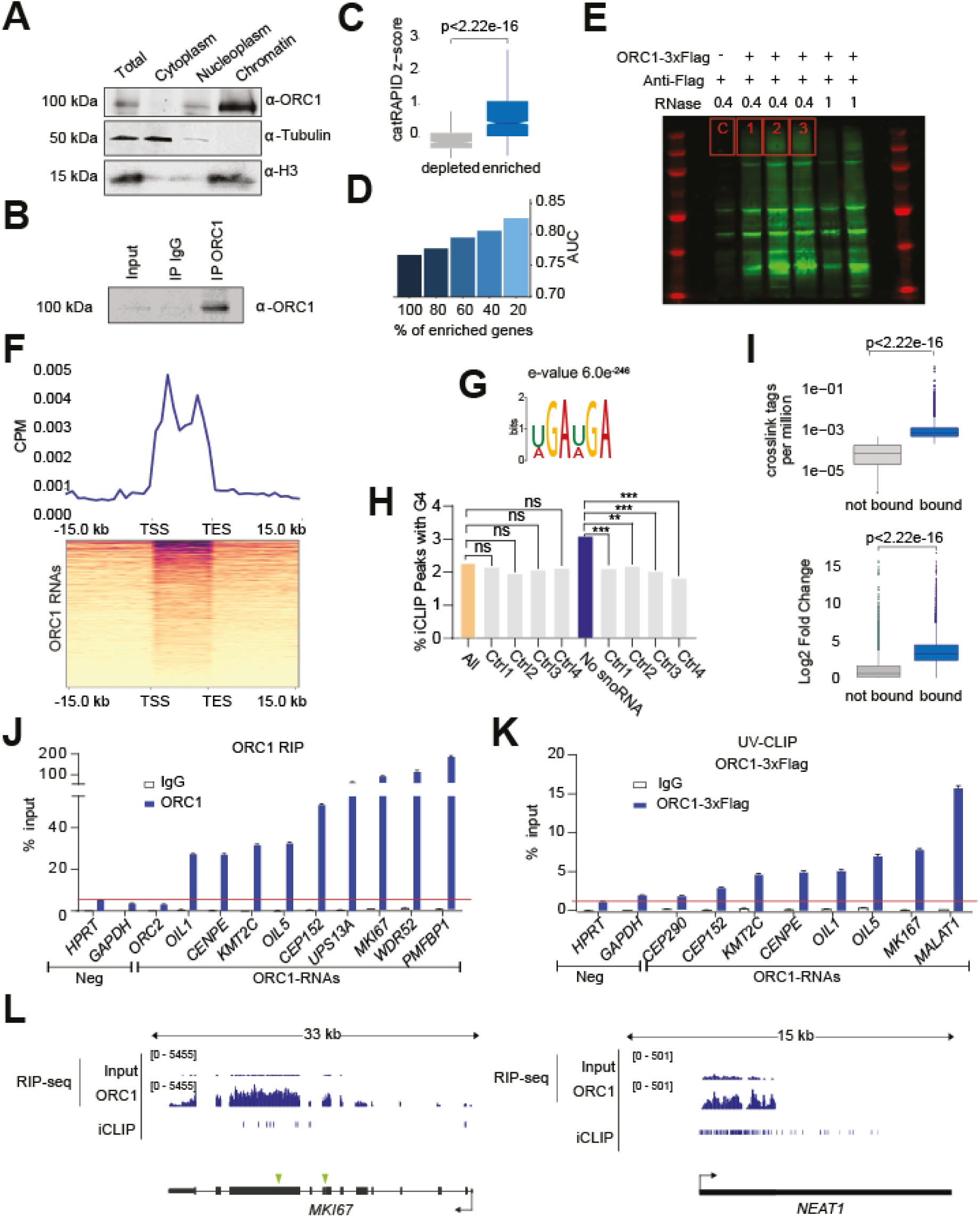
Characterization of ORC1 interactome. **(A, B)** Western blots showing **(a)** ORC1 subcellular distribution, and **(b)** ORC1 enrichment in RIP protein fraction compared to the experimental input and control IgG. **(C)** *cat*RAPID predictions of ORC1 RNA binding for depleted and enriched RNAs in ORC1 RIP-seq. **(D)** *cat*RAPID predictions of ORC1 physical interactions reporting accurate discrimination (Area Under the ROC Curve, AUC) between enriched and depleted RNAs ranked by RIP-seq fold change. **(E)** Immunoprecipitated RNA from anti- Flag iCLIP experiments, indicating ORC1-3xFlag transfection and extent of RNase digestion (units), with control (C – untransfected cells) and experimental (1, 2, 3 – ORC1-3xFlag and 0.4 RNase) gel sections analyzed marked in red. **(F)** Metagene showing ORC1 iCLIP crosslinks at extended ORC1-RNA entire genes. **(G)** MEME motif from RNA windows with ORC1 iCLIP peaks. **(H)** G4 content of RNA regions around ORC1 iCLIP peaks compared to control regions, with or without snoRNAs. **(I)** ORC1 iCLIP crosslink content (top) and RIP-seq enrichment level (bottom) of transcripts with high or low ORC1 iCLIP-defined binding**. (J, K)** ORC1-RNA RT-qPCR enrichment in **(j)** native or **(k)** UV-crosslinked RIPs (Control IgG, anti-ORC1 or anti-Flag antibodies) from a representative experiment. **(L)** Browser capture at *MKI67* or *NEAT1* ORC1-RNAs loci, presenting ORC1 RIP-seq enrichment and iCLIP peaks in HCT116 cells. Green arrows indicate positions of GAA repeats.

**Figure S3.**
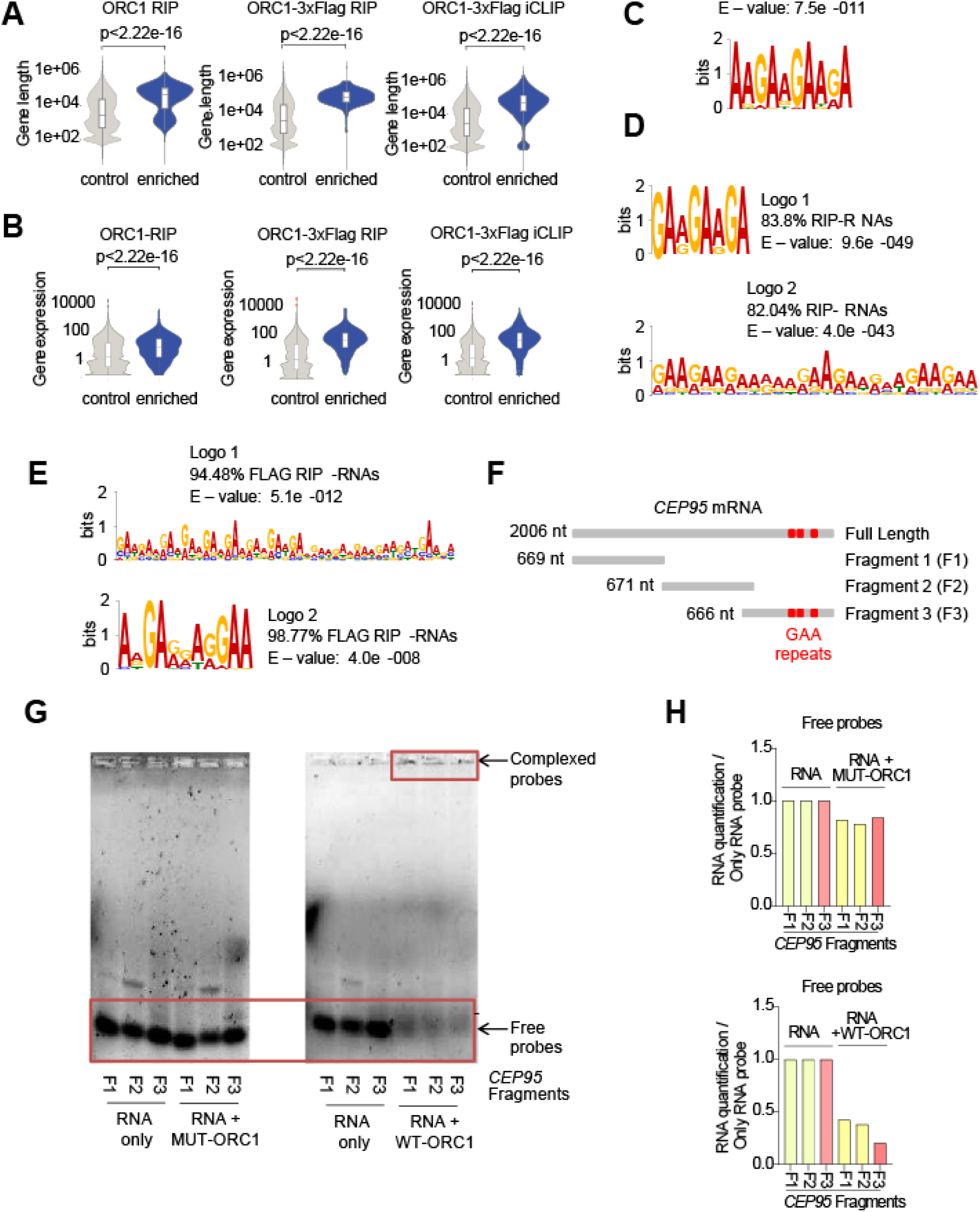
Genic features and GAA enrichment in RNAs bound to ORC1. **(A, B)** Gene **(a)** length and **(b)** expression levels of RNAs enriched by either ORC1 RIP- seq, ORC1-Flag RIP-seq, or ORC1-Flag iCLIP, compared to control genes. **(C, D, E)** MEME motifs from **(c)** entire sequences of high confidence (HC) ORC1-RNAs, **(d)** transcripts identified by ORC1 RIP-seq, or **(e)** ORC1-3xFlag RIP-seq. **(F)** Schematic of full length *CEP95* mRNA and *in vitro* synthetized RNA fragments, with GAA repeats shown in red. **(G, H) (g)** RNA-stained gels and **(h)** quantification of EMSA assays with WT and MUT versions of ORC1 RNA-binding domains (amino acids 413-511 – shown in Figure 3B) (2.5 µM), with *CEP95* RNA fragments (2.5 µM) shown in *Figure S3F*.

**Figure S4.**
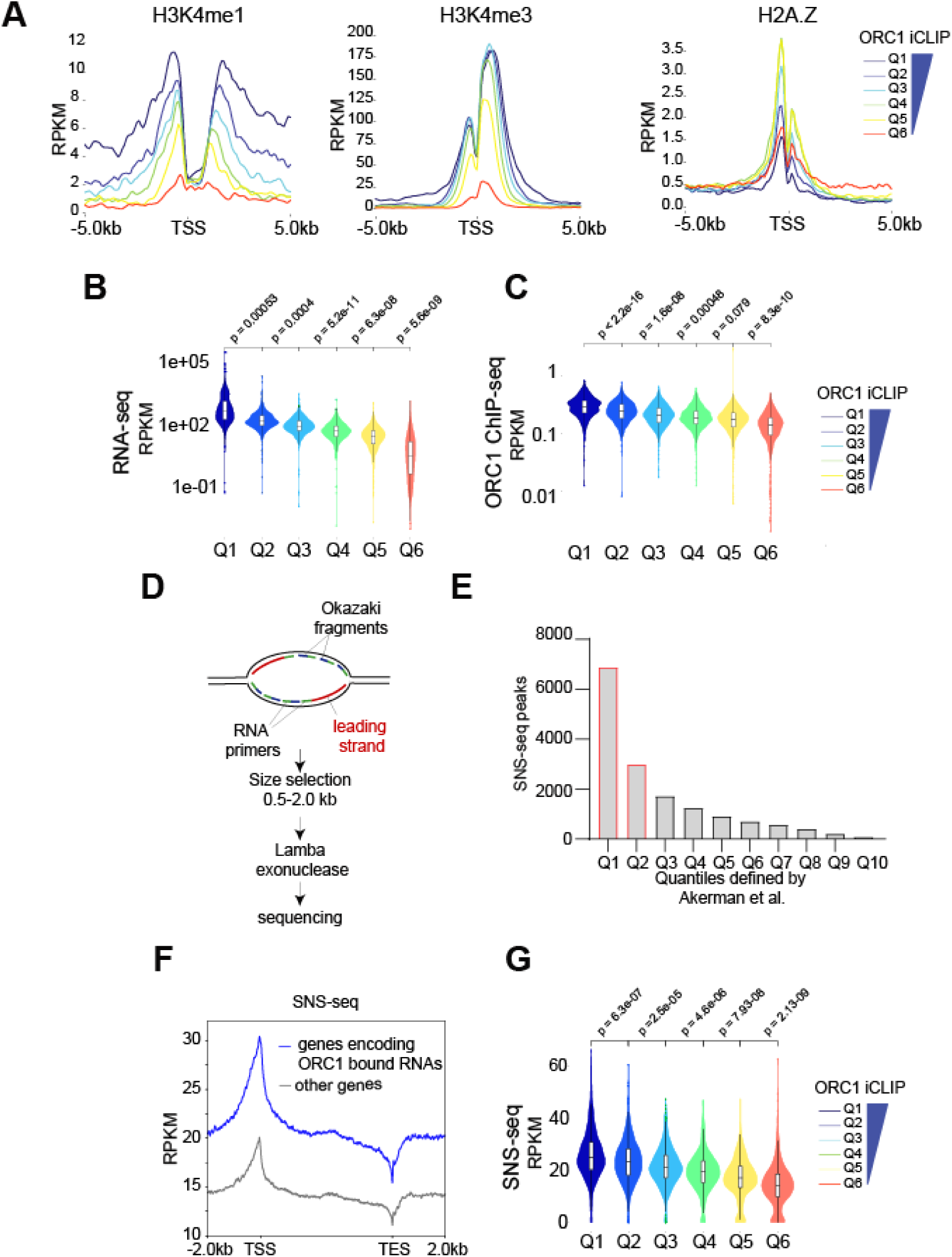
RNAs bound by ORC1 are transcribed from genes with active origins at their TSSs. **(A)** Density plots of H3K4me1, H3K4me3 and H2.AZ ChIP-seq normalized reads across six ORC1 iCLIP-defined quantiles (Q) of ORC1-RNA genes – color legend shown on the right–, centered around their TSSs (-/+ 5 kb). **(B, C)** Normalized **(b)** RNA-seq and (**c**) ORC1 ChIP-seq reads at TSSs of iCLIP-defined quantiles (Q - color legend on the right), showing *t*-test *p-values* between these groups of genes. **(D)** Schematic representation of SNS-seq protocol. Two replication forks emanate bidirectionally from origins to replicate DNA in RNA-primed leading or lagging strands. Leading strands are selected by size and λ-exonuclease digestion and sequenced. **(E)** Number of SNS-seq peaks in untreated HCT116 cells across previously defined SNS peak quantiles (Q) (Akerman et al., 2020). **(F)** Density of normalized SNS-seq reads along bodies of ORC1-RNA or control genes. **(G)** Normalized HCT116 SNS-seq reads at TSSs of iCLIP-defined gene quantiles (Q - color legend on the right), and *t*-test *p-values*.

**Figure S5.**
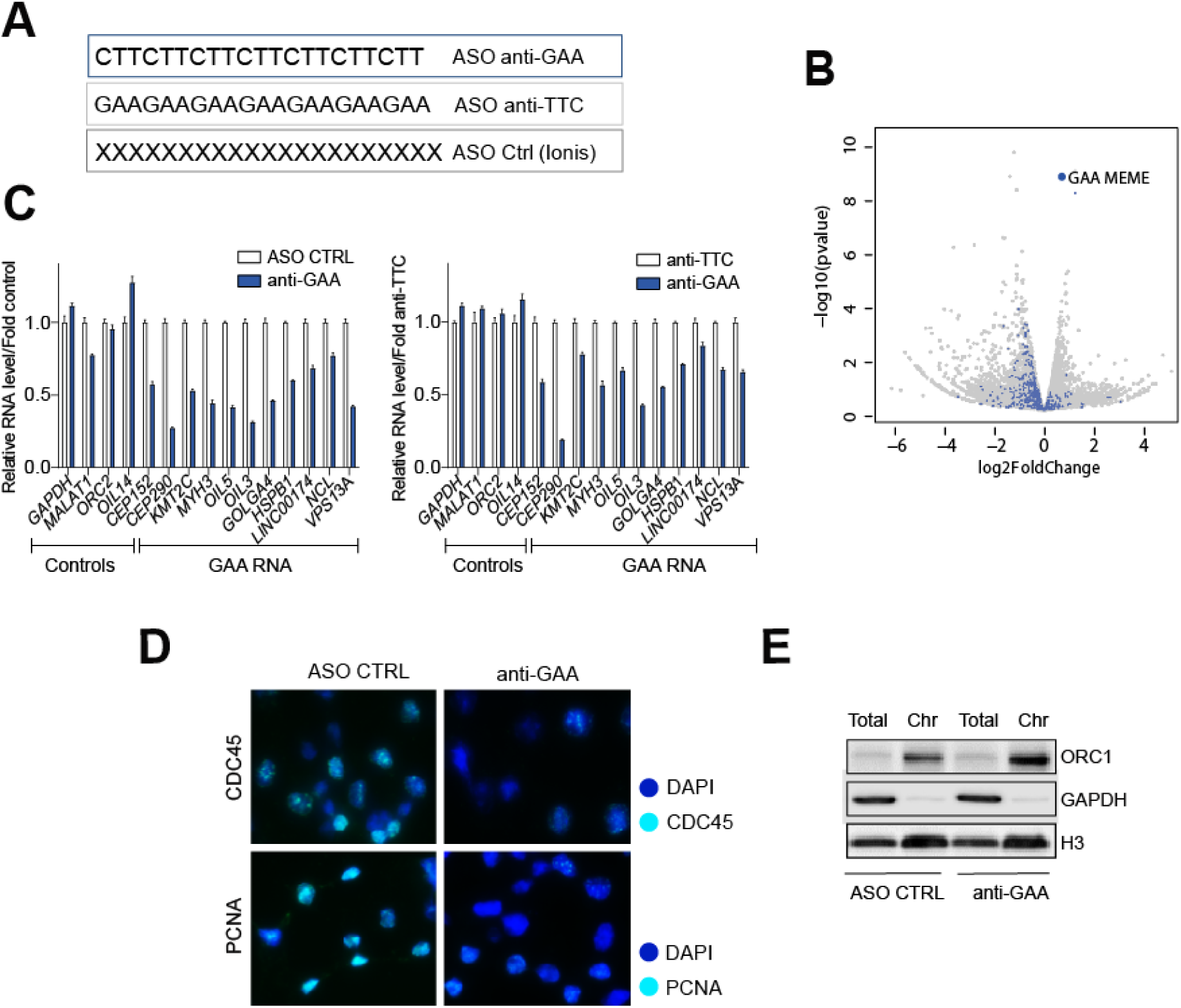
ASOs downregulate GAA-RNAs, impacting origin firing. **(A)** Design of ASOs targeting GAA repeats (anti-GAA), the reverse complement sequence (anti-TTC), and a non-targeting control. **(B)** Volcano plot of differential expression analysis (RNA-seq) in anti-GAA vs ASO control treated HCT116 cells. RNAs containing GAA sequences bound by ORC1 are shown in blue. **(C)** Representative relative RNA levels of control or ORC1-RNAs containing GAA repeats, in HCT116 cells treated with anti-GAA ASOs, relative to cells treated with control (left) or anti-TTC (right) ASOs. **(D)** Representative CDC45 and PCNA chromatin immunofluorescence images of HCT116 cells upon ASO treatments, after soluble protein washout. **(E)** Representative western blot showing ORC1 in fractioned extracts of ASO-transfected HCT116 cells.

**Figure S6.**
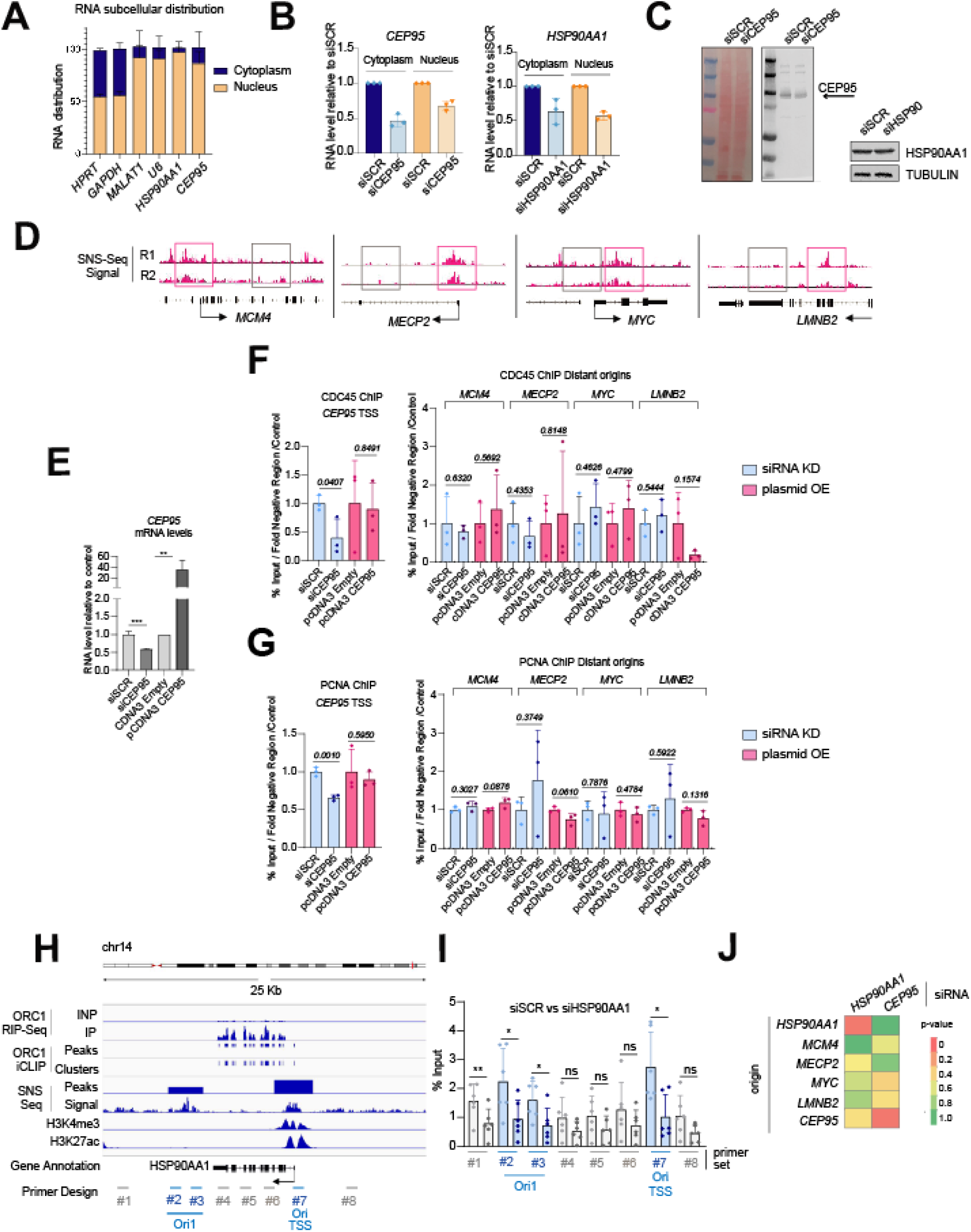
Knockdown of ORC1-RNAs reduces origin firing in *cis*. **(A)** Subcellular distribution of RT-qPCR amplified RNAs from nuclear and cytoplasmic cellular fractions of HCT116 cells. **(B, C)** *CEP95* and *HSP90AA*1 **(b)** mRNA silencing level (RT-qPCR) in HCT116 cellular fractions after 24 hours siRNAs transfection (n=3), (**c**) not affecting total protein levels. **(D)** Browser snapshots of two replicates (R) of SNS-seq data in untreated HCT116 cells, at control genomic positions of *MCM4*, *MECP2*, *MYC*, and *LMNB2* genes. Pink squares indicate SNS-enriched regions compared to proximal control gray squares. **(E)** *CEP95* mRNA levels (RT-qPCR) in knockdown (siSCR or siCEP95) and overexpression (pcDNA3 empty or CEP95) conditions, relative to controls (n=3). **(F, G)** Normalized DNA enrichment by **(f)** CDC45 and **(g)** PCNA ChIP- qPCR, at local (left) or distant (right) replication origins in conditions presented in *Figure S6E* (n=3). **(H)** Browser snapshot at *HSP90AA1* locus, showing ORC1 RIP-seq and iCLIP signals, and SNS-seq peaks and normalized reads in untreated HCT116 cells, together with public data of H3K4me3 and H3K27ac ChIP-seq. Paired oligonucleotides (#) target significant replication origins or flanking regions. **(I)** qPCR enrichment of nascent strands with primers shown in *Figure S6H*, in siSCR or siHSP90AA1 treated HCT116 cells (n=6). **(J)** Heatmap representation of *p*-*values* evaluating the statistical differences in nascent strand enrichments between siSCR and *CEP95* or *HSP90AA1* knockdowns at the indicated origins (n>5).

**Figure S7.**
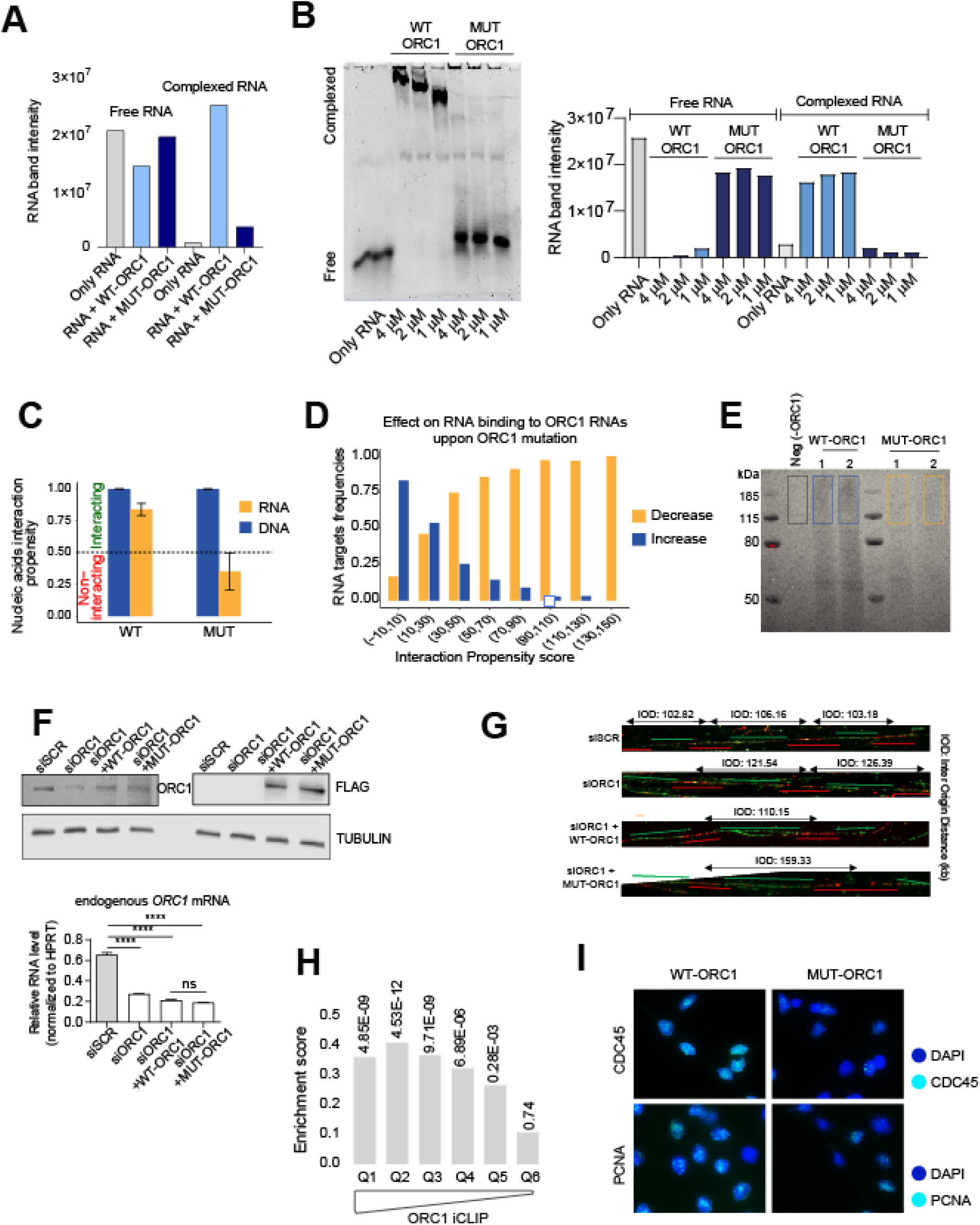
MUT-ORC1, deficient in RNA-binding, decreases origin firing. **(A)** Quantification of stained free or complexed RNA, in EMSA assays shown in Figure 4B. **(B)** RNA staining and quantification of EMSA assays, with increasing concentrations of WT/MUT ORC1 proteins (shown in Figure 4B), incubated with fragmented cellular RNA from HCT116 cells (2 µM) **(C)** *CleverSuite* predicted DNA and RNA binding activity of wild type (WT) and mutant (MUT) ORC1. **(D)** *cat*RAPID omics v2 prediction of the increase or decrease of the protein-RNA interaction propensity, caused by the mutation of ORC1 and different bins of ORC1-RNAs. **(E)** Staining of immunoprecipitated RNA from control or anti-Flag iCLIP experiments, in cells transiently expressing WT or MUT- ORC1 tagged with Flag, or untransfected cells as negative control. Boxes indicate gel sections at and above ORC1-expected molecular weight. **(F)** Western blots and relative *ORC1* mRNA levels showing knockdown of endogenous ORC1 and overexpression of exogenous WT and MUT-ORC1 tagged with Flag, in rescue experiments shown in Figure 4D. **(G)** Representative labeled DNA fibers showing distances between interspersed replication origins in WT or MUT-ORC1 rescue experiments shown in Figure 4D. **(H)** Enrichment score and associated adjusted *p-values* from GSEA analysis shown in Figure 4F, for genes in individual iCLIP-defined quantiles. **(I)** Representative CDC45 and PCNA chromatin immunofluorescence images in HCT116 cells stably expressing WT or MUT-ORC1 tagged with Flag, after soluble protein washout.

**Figure S8.**
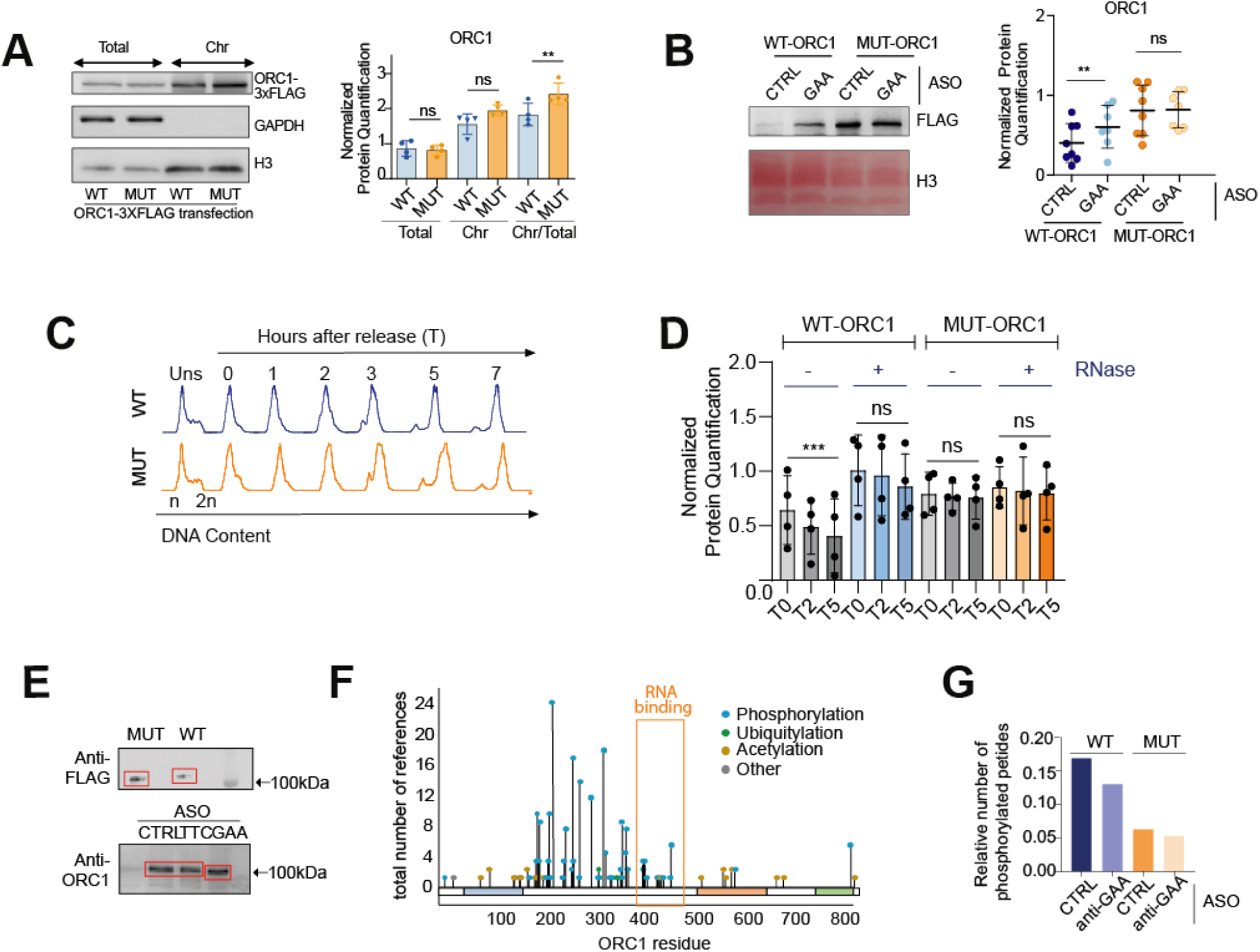
RNA-dependent ORC1 phosphorylation controls ORC1 chromatin release. **(A)** Western blot and protein quantifications of ORC1-3xFlag in total and chromatin extracts of HCT116 cells transiently expressing WT or MUT- ORC1, with calculated chromatin/total ORC1 ratio (n=4). **(B)** Representative western blot and protein quantifications of WT or MUT- ORC1-3xFlag in the chromatin fraction of control or GAA- RNA knocked down HCT116 stable cells (n=8). **(C)** Flow cytometry analysis of synchronized HCT116 cells, transfected with WT or MUT- ORC1-3xFlag, representing DNA content (n or 2n) of unsynchronized cellular populations, or at different hours (T) of thymidine release. **(D)** Quantification of western blot presented in Figure 5D, showing WT and MUT-ORC1 chromatin association in synchronized cells (T as in *Figure S8C*) +/- RNase A treatment. **(E)** Western blots showing (up) mobility shift differences between Flag-tagged WT and MUT-ORC1, and (bottom) between endogenous ORC1 in control or GAA-RNA depletion conditions, in HCT116 cells. **(F)** PhosphositePlus website representation of referenced ORC1 post-translational modifications along protein residues (x axis), highlighting its RNA-binding region. **(G)** Number of detected phosphorylated peptides of WT or MUT-ORC1 by mass spectrometry, relative to the total number of detected peptides, in control or GAA-RNA knockdown conditions.

**Figure S9.**
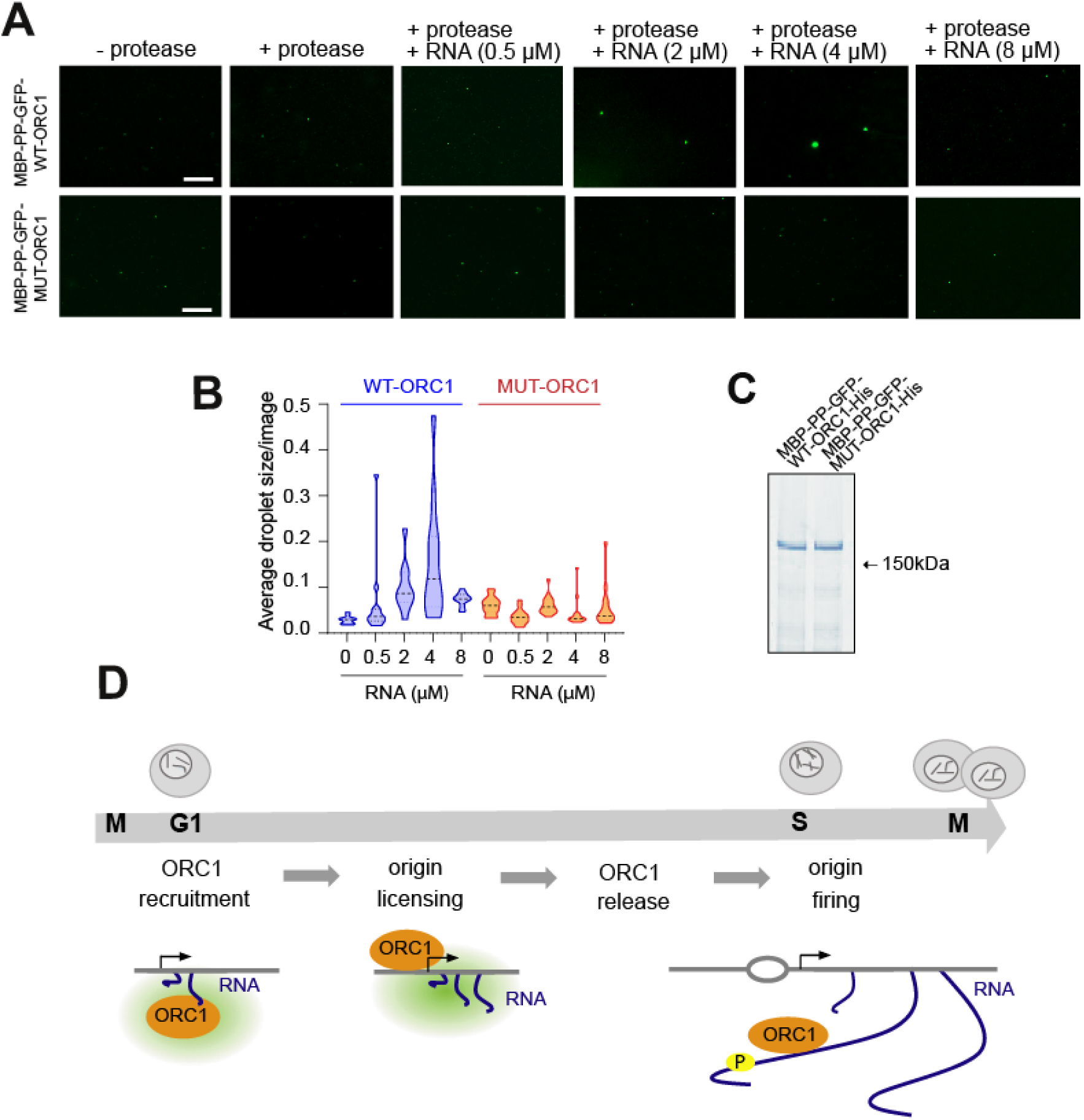
Phase separation of ORC protein on RNA. **(A)** Representative images of droplets containing MBP-PP-GFP-WT-ORC1 and MBP-PP-GFP-MUT-ORC1 proteins (shown in *Figure S9C*), treated or untreated with protease to release MBP, and incubated with the indicated concentrations of RNA oligo (Scale bars, 10µm). **(B)** Quantification of LLPS MBP-PP-GFP-WT-ORC1 or MBP-PP-GFP-MUT-ORC1 droplets as a function of RNA concentrations. Bold dashed lines indicate median; dotted lines indicate quartiles. **(C)** Coomassie blue staining of purified full-length tagged WT or MUT-ORC1. **(D)** Speculative model for RNA binding to ORC1 in different phases of the cell cycle. The green area represents liquid-liquid phase separation by ORC1 (and possibly other initiation factors) interacting with RNA.

**Table S1.** iCLIP sequencing reads.

| anti-FLAG IP | Uniquely mapped reads | Mapping reads | Multimapping reads | All reads | Unmapped reads |
| --- | --- | --- | --- | --- | --- |
| Untransfected (negative control) | 764365 | 2971915 | 2207550 | 3583696 | 611781 |
| ORC1-3xFlag expressing cells (Replicate 1) | 7319329 | 40232831 | 32913502 | 41963528 | 1730697 |
| ORC1-3xFlag expressing cells (Replicate 2) | 8333176 | 44541088 | 36207912 | 46127516 | 1586428 |
| ORC1-3xFlag expressing cells (Replicate 3) | 9928271 | 60729943 | 50801672 | 61965266 | 1235323 |

**Table S2.** List of plasmids.

| Plasmid Number | Backbone | Insert | Comment |
| --- | --- | --- | --- |
| 1 | pCDNA3.1 | WT-ORC1-3xFlag | Codon-optimized |
| 2 | pCDNA3.1 | *MUT-ORC1-3xFlag | Codon-optimized |
| 3 | pBABE | Halo-WT-ORC1 | Cloned from plasmid 1 |
| 4 | pBABE | *Halo-MUT-ORC1 | Cloned from plasmid 2 |
| 5 | pCDNA3.1 | CEP95 | Clone (OHu50782C) |
| 6 | pMALp-c2E | MBP-pp-GFP-hORC1wt-6xhis | Stillman Lab |
| 7 | pMALp-c2E | *MBP-pp-GFP-hORC1mut-6xhis |  |

\*ORC1 codons for arginines (R), in amino acid positions 441, 444 and 465, were substituted by alanine (A) codon GCC, to generate ORC1 RNA-binding mutant (R441A, R444A, R465A). Plasmids with these cDNA sequences were manually designed and synthesized by GenScript.

**Table S3.**
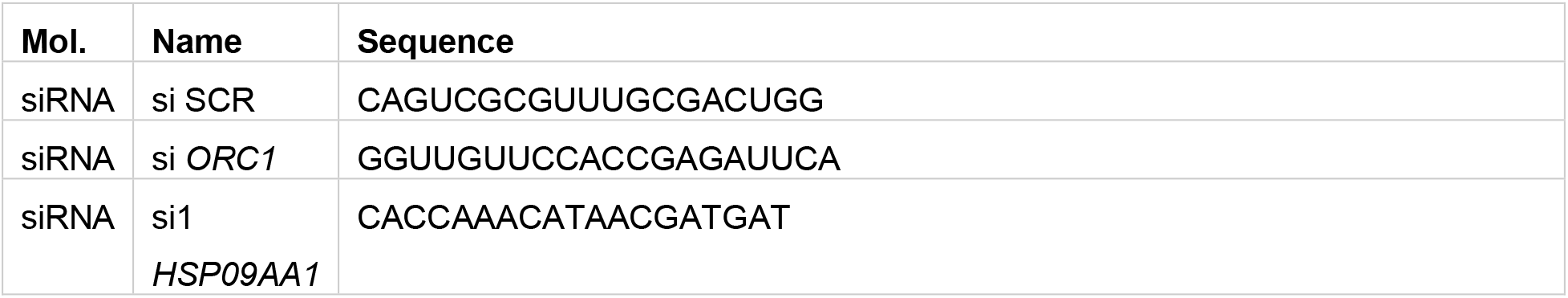
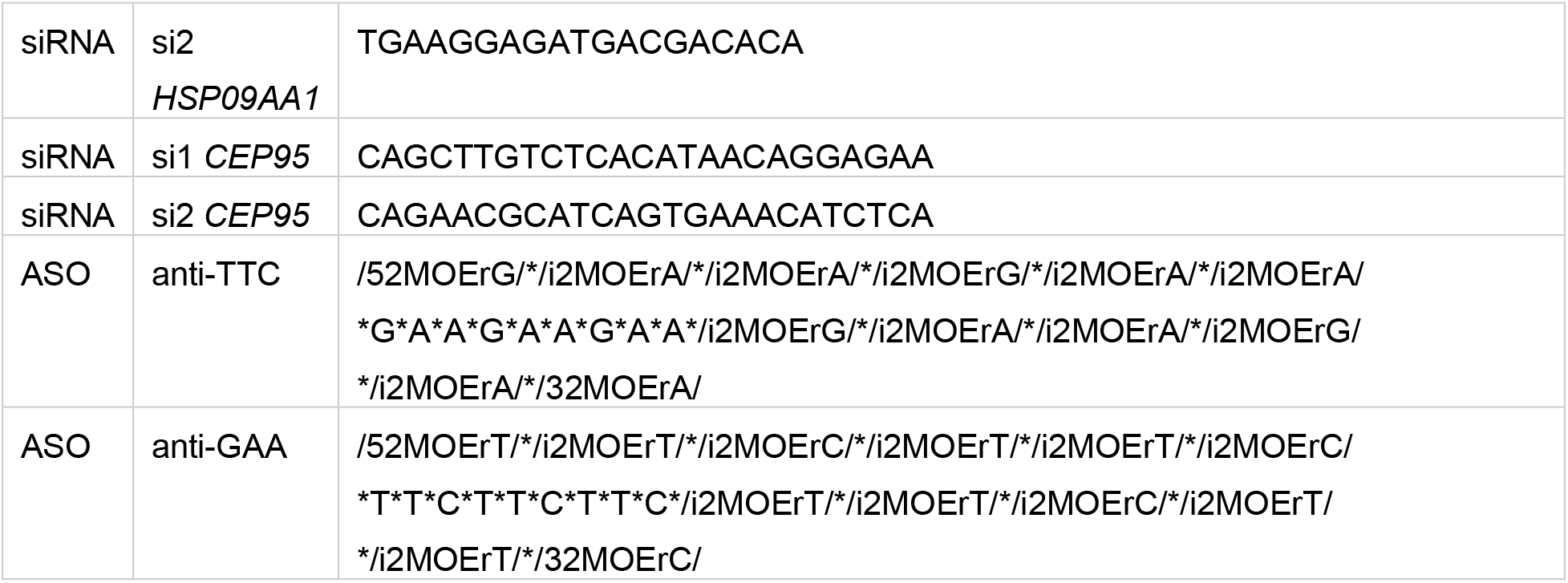
Sequence of siRNAs and ASOs.

**Table S4.**
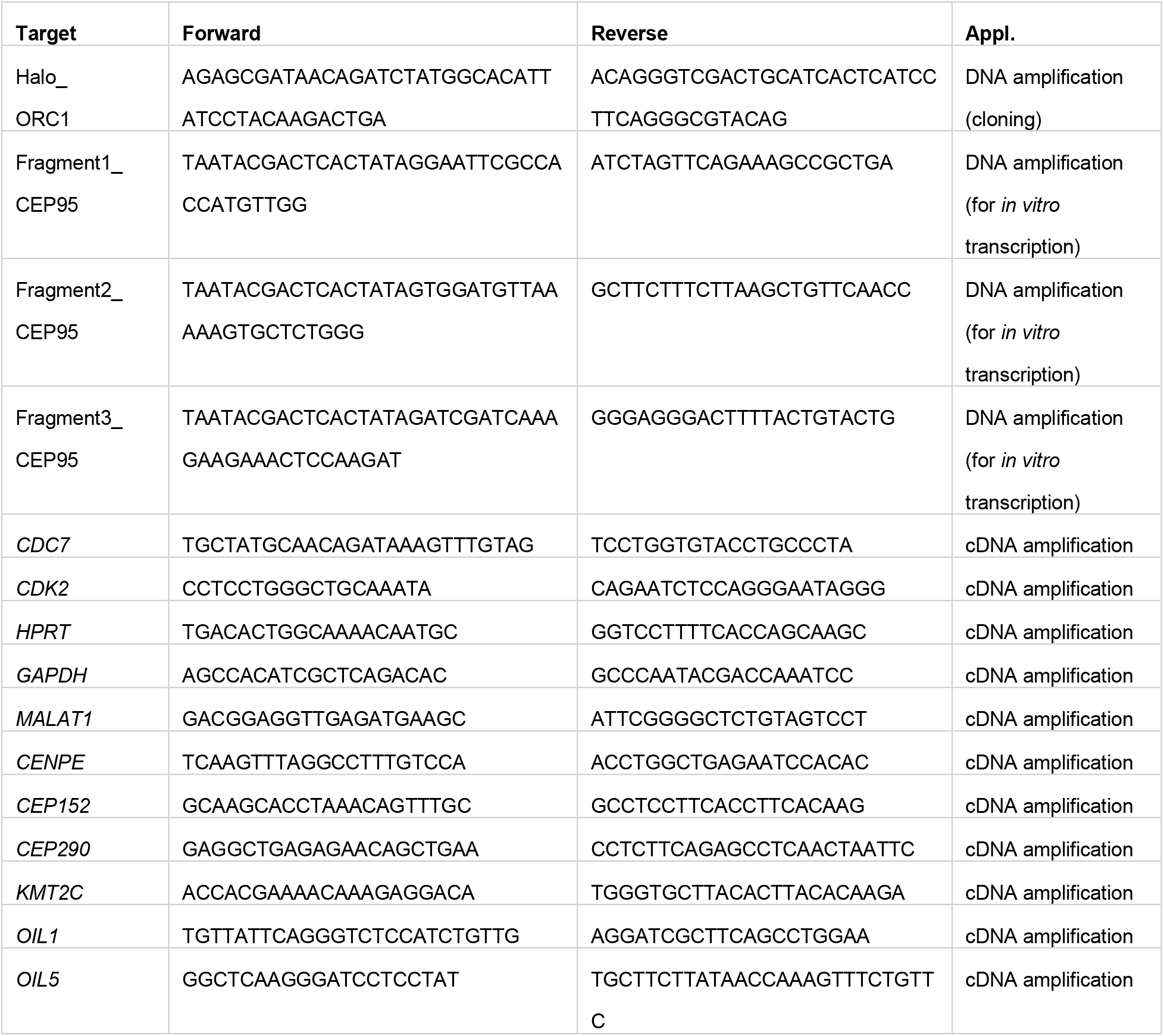
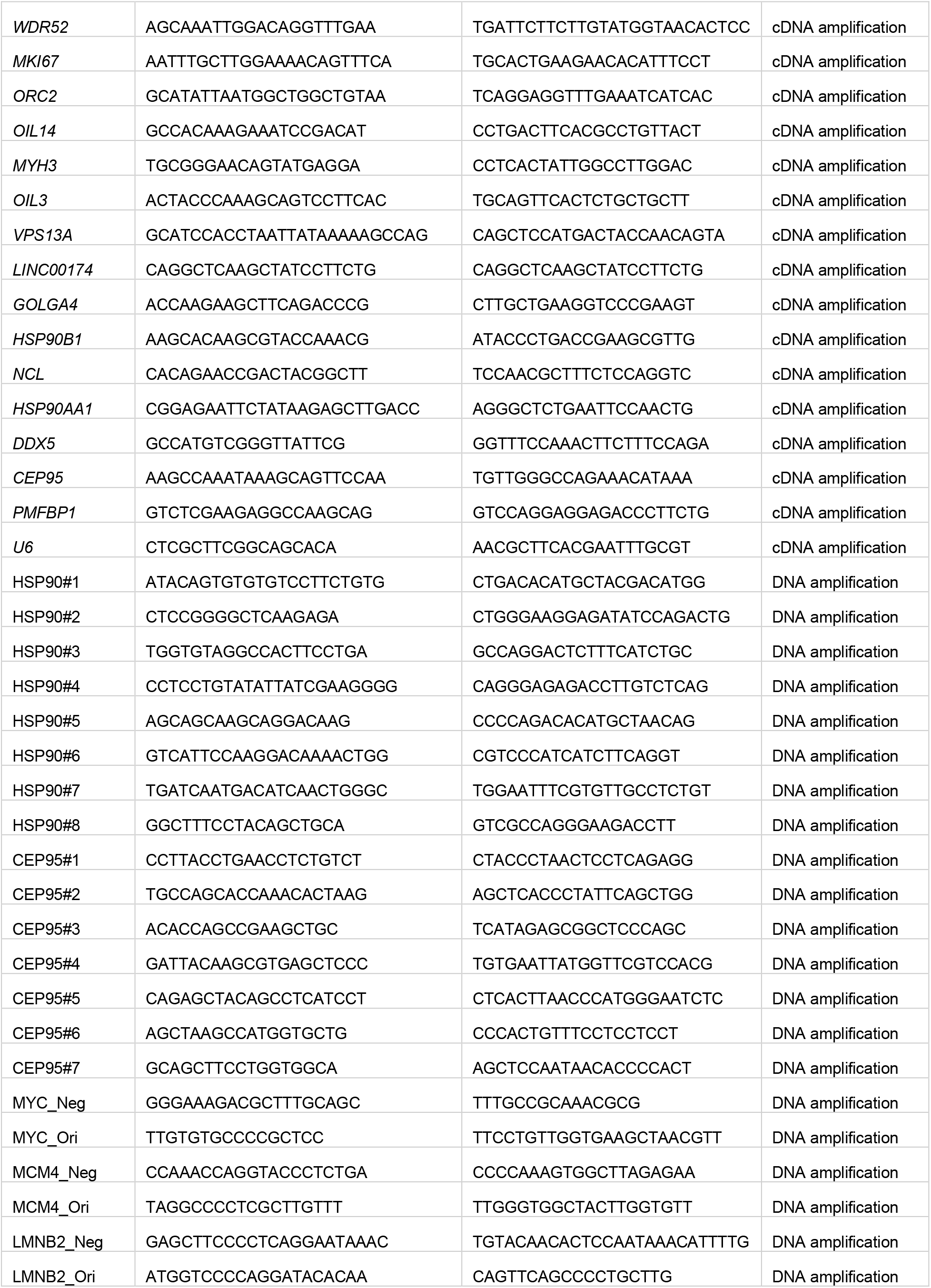
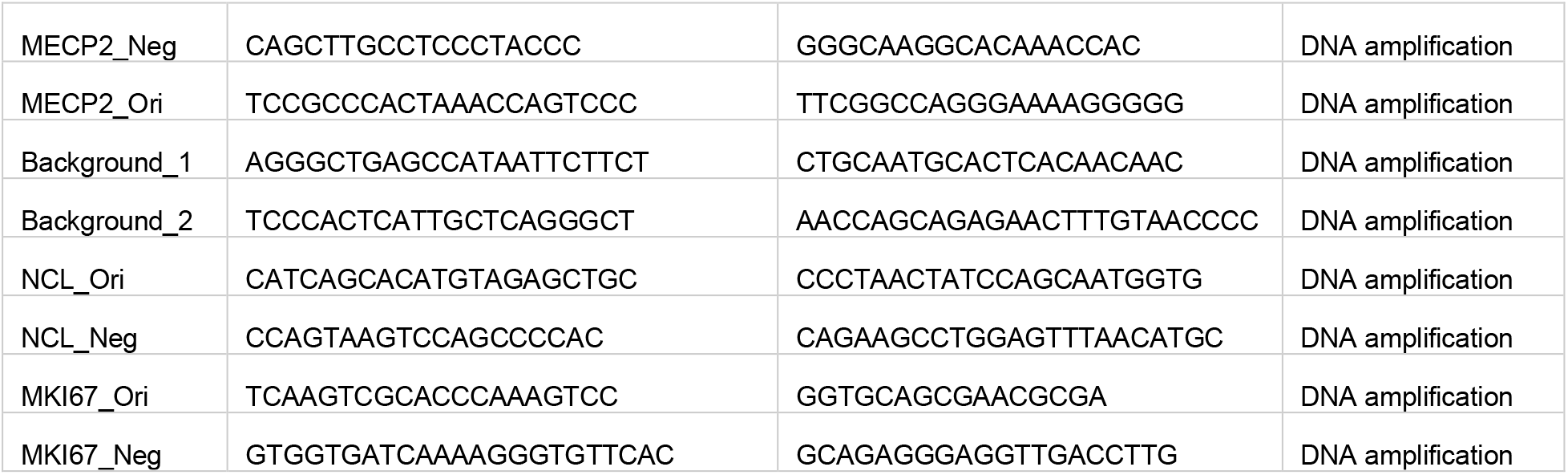
Sequence of oligonucleotides and application.

**Table S5.** List of antibodies used for western blotting.

| Primary Antibodies | Reference |
| --- | --- |
| ORC1 (F-10) | sc-398734 (Santa Cruz) |
| ORC1 (7A7) | sc-23887 (Santa Cruz) |
| GAPDH (HRP-Conj) | 3683 (Cell Signalling) |
| Tubulin | T5158 (Sigma) |
| H3 | ab1791 (Abcam) |
| Flag M2 | F3165 (Sigma) |
| PCNA (PC10) | sc-56 (Santa Cruz) |
| p53 | MABE327 (Sigma) |
| HSP90 | 4874 (Cell Signalling) |
| CEP95 | MBS3007560<br>(mYbIOsOURCE) |

**Table S6.** ORC1 orthologues. Orthologous proteins from vertebrates and metazoans are listed.

| Species | Protein ID | Database |
| --- | --- | --- |
| *Abeoforma whisleri* | g.2331 | Multicellgenome |
| *Alligator mississippiensis* | A0A151M4M2 | Uniprot |
| *Amphimedon queenslandica* | A0A1X7VWI9 | Uniprot |
| *Anolis carolinensis* | G1KHI9 | Uniprot |
| *Apis mellifera* | A0A088AEY1 | Uniprot |
| *Arabidopsis thaliana* | Q710E8 / Q9SU24 | Uniprot |
| *Branchiostoma floridae* | C3ZFR4 | Uniprot |
| *Caenorhabditis elegans* | Q9XX17 | Uniprot |
| *Callorhinchus milii* | V9KDF9 | Uniprot |
| *Capsaspora owczarzaki* | A0A0D2WSH5 | Uniprot |
| *Chelonia mydas* | M7B8T8 | Uniprot |
| *Crassostrea gigas* | K1Q150 | Uniprot |
| *Danio rerio* | A0A0R4IFH7 | Uniprot |
| *Daphnia pulex* | E9G6A6 | Uniprot |
| *Drosophila melanogaster* | O16810 | Uniprot |
| *Gallus gallus* | Q5ZMC5 | Uniprot |
| *Helobdella robusta* | T1EDE2 | Uniprot |
| *Homo sapiens* | Q13415 | Uniprot |
| *Ixodes scapularis* | B7Q2L8 | Uniprot |
| *Latimeria chalumnae* | H3AUF8 | Uniprot |
| *Lepisosteus oculatus* | W5MGU6 | Uniprot |
| *Lottia gigantea* | V4BTT9 | Uniprot |
| *Micromonas pusilla* | C1N5G9 | Uniprot |
| *Mnemiopsis leidyi* | ML02754a | NHGRI |
| *Monodelphis domestica* | F6QFY9 | Uniprot |
| *Monosiga brevicollis* | A9V9J0 | Uniprot |
| *Mus musculus* | Q9Z1N2 | Uniprot |
| *Nematostella vectensis* | A7STK3 | Uniprot |
| *Octopus bimaculoides* | A0A0L8FWE5 | Uniprot |
| *Oikopleura dioica* | E4XD69 | Uniprot |
| *Ornithorhynchus anatinus* | F6TS55 | Uniprot |
| *Oryzias latipes* | H2MEI0 | Uniprot |
| *Oscarella carmela* | m.102310 | Compagen |
| *Physcomitrium patens* | A9SL85 | Uniprot |
| *Pirum gemmata* | g.18948 | Multicellgenome |
| *Rhizophagus irregularis* | U9T192 | Uniprot |
| *Sphaeroforma arctica JP610* | A0A0L0GEU8 | Uniprot |
| *Saccharomyces cerevisiae* | P54784 | Uniprot |
| *Sycon ciliatum* | scpid68717 | Compagen |
| *Saccoglossus kowalevskii* | XP_002738319 | NCBI RefSeq |
| *Strigamia maritima* | T1JIW5 | Uniprot |
| *Schizosaccharomyces pombe* | P54789 | Uniprot |
| *Salpingoeca rosetta* | F2UK60 | Uniprot |
| *Trichoplax adhaerens* | B3S114 | Uniprot |
| *Tribolium castaneum* | D6WQP2 | Uniprot |
| *Taeniopygia guttata* | H0ZF97 | Uniprot |
| *Xenopus tropicalis* | F6SND7 | Uniprot |

